# Brain and blood single-cell transcriptomics in acute and subacute phases after experimental stroke

**DOI:** 10.1101/2023.03.31.535150

**Authors:** Lidia Garcia-Bonilla, Ziasmin Shahanoor, Rose Sciortino, Omina Nazarzoda, Gianfranco Racchumi, Costantino Iadecola, Josef Anrather

**Affiliations:** The Feil Family Brain and Mind Research Institute, Weill Cornell Medicine, New York, NY 10021

## Abstract

Cerebral ischemia triggers a powerful inflammatory reaction involving both peripheral leukocytes and brain resident cells. Recent evidence indicates that their differentiation into a variety of functional phenotypes contributes to both tissue injury and repair. However, the temporal dynamics and diversity of post-stroke immune cell subsets remain poorly understood. To address these limitations, we performed a longitudinal single-cell transcriptomic study of both brain and mouse blood to obtain a composite picture of brain-infiltrating leukocytes, circulating leukocytes, microglia and endothelium diversity over the ischemic/reperfusion time. Brain cells and blood leukocytes isolated from mice 2 or 14 days after transient middle cerebral artery occlusion or sham surgery were purified by FACS sorting and processed for droplet-based single-cell transcriptomics. The analysis revealed a strong divergence of post-ischemic microglia, macrophages, and neutrophils over time, while such diversity was less evident in dendritic cells, B, T and NK cells. Conversely, brain endothelial cells and brain associated-macrophages showed altered transcriptomic signatures at 2 days post-stroke, but low divergence from sham at day 14. Pseudotime trajectory inference predicted the in-situ longitudinal progression of monocyte-derived macrophages from their blood precursors into day 2 and day 14 phenotypes, while microglia phenotypes at these two time points were not connected. In contrast to monocyte-derived macrophages, neutrophils were predicted to be continuously de-novo recruited from the blood. Brain single-cell transcriptomics from both female and male aged mice did not show major changes in respect to young mice, but aged and young brains differed in their immune cell composition. Furthermore, blood leukocyte analysis also revealed altered transcriptomes after stroke. However, brain-infiltrating leukocytes displayed higher transcriptomic divergence than their circulating counterparts, indicating that phenotypic diversification into cellular subsets occurs within the brain in the early and the recovery phase of ischemic stroke. In addition, this resource report contains a searchable database https://anratherlab.shinyapps.io/strokevis/ to allow user-friendly access to our data. The StrokeVis tool constitutes a comprehensive gene expression atlas that can be interrogated at the gene and cell type level to explore the transcriptional changes of endothelial and immune cell subsets from mouse brain and blood after stroke.

## Introduction

Ischemic stroke (IS) accounts for about 87% of all strokes and occurs when a vessel supplying blood to the brain is obstructed. Approaches for treatment of acute stroke are targeted at rapid recanalizing of the blocked artery by endovascular thrombectomy and/or administration of thrombolytic agents ^1,2^. Despite recent successes in both prevention and treatment, stroke is yet the second leading cause of death and third leading cause of disability worldwide ^3,4^.

The immune system actively participates in the acute and chronic pathogenesis of IS. Damaged neurons lead to a secondary inflammatory reaction that aggravates brain injury, increasing neurologic deficits ^5-7^. This response progresses for days to weeks and involves glial and brain endothelium activation, recruitment of peripheral immune cells, and release of cytokines. Whereas there is evidence that the acute inflammatory response contributes to the progression of ischemic brain injury, more recent research points at a more multifaceted role of immune cells in brain ischemia, where they participate in repair processes during the sub-acute and chronic stages ^8-11^. In addition, post-stroke chronic inflammation and adaptive immunity have been implicated in the long-term sequelae such as depression and dementia ^12^.

Emerging single-cell RNA sequencing (scRNA-seq) studies reveal a highly transcriptional cellular heterogeneity in response to IS ^13^, supporting the previously reported functional plasticity of immune cells after stroke ^14^. Although the precise mechanisms inducing cellular diversity have not yet been determined, signals released by damaged brain cells as well as other environmental cues like metabolic alterations after ischemia, are most likely to induce phenotypic differentiation of the brain cells. Furthermore, the ischemic brain and systemic immunity interact in a bidirectional fashion. While the immune system supplies the brain with immune cells that participate in the local inflammatory response, neural and humoral factors generated by the ischemic brain communicate to peripheral organs ^7^. Thus, activation of the immune system through brain-derived molecules or via autonomic nervous system, might lead to transcriptional differentiation of immune cells before entering the brain ^7,15^

Some studies using scRNA-seq technology have begun to decipher the cellular heterogeneity of brain cells after IS ^16-20^. Here we used single-cell RNA sequencing to gain deeper insights into the impact of IS on transcriptional diversity of brain immune cells, brain endothelial cells (EC) and peripheral blood leukocytes. Specifically, this study focuses on assessing differences of transcriptomic signatures related to cell origin (brain resident vs. recruited cells), cellular localization (periphery vs. brain), time post-stroke (acute versus sub-acute) and age (young vs. aged mice).

## Results

### Dynamics of mouse brain and blood cell heterogeneity over the ischemic-reperfusion time

To obtain a comprehensive picture of the immune cell diversity and temporal dynamics over the acute and subacute phases after cerebral ischemia, we isolated immune cells from the brains of stroke mice and performed droplet-based scRNA-seq. We also isolated endothelial cells (EC), the first cells to interact with leukocytes migrating into the ischemic tissue. We prepared brain single cell suspensions from mice 2 days after sham surgery (Sham) or 2 and 14 days after stroke (D02 and D14, respectively, Fig.1A). Subsequently, we flow sorted peripheral immune cells and border-associated macrophages (BAM; CD45^hi^), microglia (CD45^int^CX3CR1^+^), and EC (CD45^low^Ly6C^hi^)^21,22^. Cells were mixed in a 6:1:1 ratio (CD45^hi^:microglia:EC) and processed for droplet-based RNA library preparation (Fig.1A). We enriched the samples for CD45^hi^ cells because we anticipated higher cellular heterogeneity of CD45^hi^ population than in either microglia or EC. We detected 13 distinct major cell type clusters in 43269 cells after performing unsupervised clustering and Uniform Manifold Approximation and Projection (UMAP). On the basis of the expression of established marker genes and unsupervised cell type annotation, clusters were identified as microglia (Mg), BAM, monocyte-derived cells (MdC), granulocytes (Gran), mast cells (MaC), dendritic cells (DC), T cells (Tc), NK cells (NK), B cells (Bc), endothelial cells (EC), vascular mural cells (MC), epithelial-like cells (Epi), and oligodendrocytes (OD) (Fig.1B; Fig. S3). As anticipated, we observed that the relative abundance of each cell cluster changed notably with ischemia-reperfusion. For instance, MdC increased at D02, whereas the increase in lymphocytes (Tc and NK) was more pronounced at D14, as previously reported ^6,23,24^. Amid brain resident cells, we observed divergent clustering of microglia across Sham, D02 and D14, indicating that ischemic injury markedly modified microglia transcriptomes. On the other hand, the UMAP representation indicated that BAM and EC clusters were less heterogeneous across groups (Fig. 1B).

**Figure 1.**
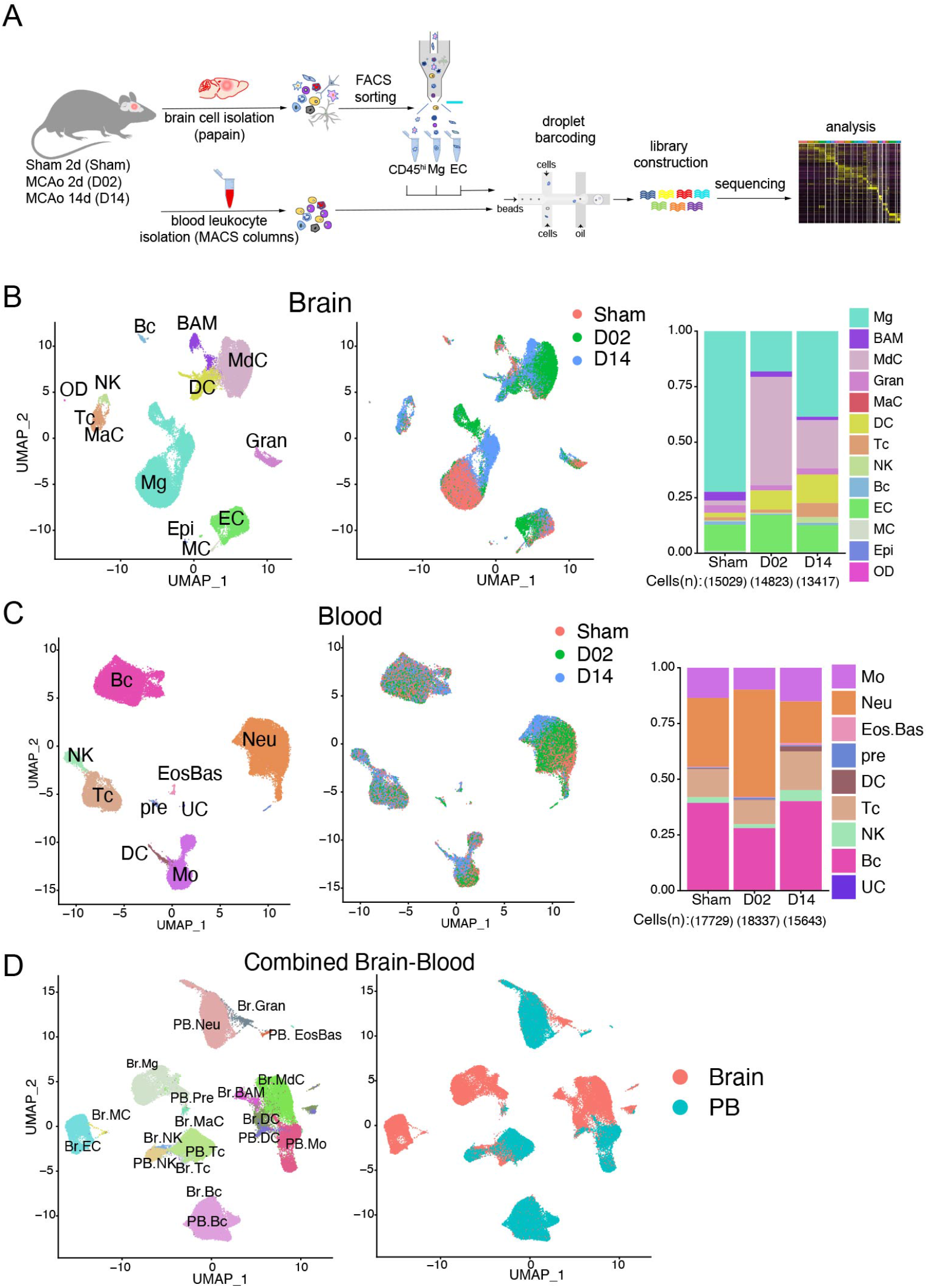
Single-cell transcriptomic profiling of mouse brain and blood cells after transient focal cerebral ischemia. **(A)** Schematic representation of Drop-Seq scRNA-seq pipeline used to analyze brain and blood cells isolated from either control surgery (Sham) or stroke mice 2 and 14 days (D02, D14) after injury. Brain cells were dissociated by enzymatic digestion with papain. Infiltrating leukocytes (CD45hi), microglia (Mg) and endothelial cells (EC) were isolated by flow cytometry sorting. Blood leukocytes were purified after erythrocyte removal. Brain and blood single cell suspensions were subjected to Drop-Seq, sequencing and analysis. **(B, C)** *Left*: Uniform Manifold Approximation and Projection (UMAP) plot representing color-coded cell clusters identified in merged brain (B) or blood (C) single-cell transcriptomes; *Middle*: UMAP of 3 color-coded time point overlay of brain (B) or blood (C) single-cell transcriptomes; *Right*: bar graph showing relative frequencies of each cell type across Sham, D02 and D14 groups of either brain (B) or blood (C) identified cell type clusters. **(D)** *Left*: UMAP plot of the combined brain (Br) and blood (PB) dataset showing cell clustering similarities between brain and blood Gran, Tc, Bc and brain myeloid cells (BAM, MdC, DC) with blood monocytes (left). *Left*: UMAP plot shows the distribution of brain and blood cells overlying the identified clusters. Border-associated macrophages (BAM), monocyte-derived cells (MdC), granulocytes (Gran), mast cells (MaC), dendritic cells (DC), T cells (Tc), NK cells (NK), B cells (Bc), vascular mural cells (MC), epithelial-like cells (Epi), oligodendrocytes (OD); Eosinophils-Basophils (EosBas); Monocytes (Mo); hematopoietic precursors (pre); unclassified (UC).

Because brain ischemia elicits a systemic immune response that ultimately affects stroke outcome ^7^, we next sought to explore the transcriptomes of peripheral blood cells from stroke mice. We performed scRNA-seq of purified blood leukocytes from Sham, D02 and D14 mice. Leukocytes grouped into 8 distinct clusters that were annotated by the expression of commonly used marker genes, as follows: monocytes (Mo), granulocytes (Gran), eosinophils/basophils (Eos/Bas), dendritic cells (DC), T cells (Tc), NK cells (NK), B cells (Bc), various precursors (pre), and one unclassified cluster (UC) (Fig.1C; Fig. S4). Overall, the major peripheral blood immune cell types showed conserved positioning in the UMAP space across Sham, D02 and D14 groups. Only Gran and Mo clusters were slightly divergent at D14 from their respective Sham and D02 clusters. Thus, these results point to lesser phenotypic differentiation in circulating leukocytes than in brain recruited leukocytes after stroke.

To assess transcriptional similarities between leukocytes from both tissues we analyzed a combined brain and blood dataset. We observed that non-brain resident immune cells clustered together independently of their tissue origin. On the other hand, whereas BAM, MdC, and brain DC clustered with blood monocytes and DC, the microglia cluster remained spatially separated from blood myeloid cells reflecting their unique transcriptome (Fig. 1D).

### Cerebral ischemia induces different transcriptomic states in microglia at early and late phases

To determine whether distinct transcriptional programs of microglia develop over the ischemia-reperfusion time, we re-clustered microglia which resulted in 8 clusters (Mg1-8) all showing high expression of canonical microglial markers *Hexb*, *Olfml3*, and *Fcrls* (Fig. 2A-D; Fig. S5A). Sham microglia mostly fell into clusters Mg1 and Mg2, which exhibited high expression of genes found in homeostatic microglia including *P2ry12*, *Tmem119*, and *Sall1*. Whereas Mg1 was mainly characterized by expression of homeostatic genes (*Siglech*, *P2ry12*, *Tmem119*), Mg2 showed upregulation of immediate early genes (*Jun*, *Fos*, *Erg1, Klf2*, *Klf4* and *Atf3*), a group of transcription factors typically expressed in adult microglia and involved in establishing microglia surveilling functions ^25-28^. In line with recent reports, we observed that microglial homeostatic genes, such as *P2ry12* and *Tmem119*, were downregulated after cerebral ischemia ^16,29^ (Fig. 2B). The expression of *Apoe, Lpl, Spp1, Clec7a* or *Cst7* genes, which have been linked to microglial responses to demyelination ^30^, Alzheimer’s disease ^31^, as well as stroke ^16,29^, were upregulated in both D02 and D14 microglia (Fig. 2A-D). Furthermore, cerebral ischemia led to distinct D02 and D14 microglia clusters. D02 microglia was mainly composed of Mg4 and Mg5 clusters, while the most prominent clusters at D14 were Mg3, Mg6, and Mg7 (Fig. 2C). Among D02 microglia clusters, Mg4 showed high expression of genes related to the clearance of damaged cells and tissue repair *(Spp1*, *Msr1* and *Lgals3)* ^29,32-34^, and high expression of chemokine genes (*Ccl2*, *Ccl12*) ^35^. The Mg5 cluster was characterized by the expression of mitotic genes including *Top2A, Mki67* and *Stmn1* ^20^, indicating the presence of proliferative microglia during the acute phase of stroke. Mg3, the predominant cluster at D14, was mainly defined by expression of genes associated with disease-associated microglia (DAM) including *Apoe, Cst7, Clec7a, Lyz2, Lgals3bp, Igf1*, and *Lpl* ^31^ (Fig. S5B). Similar to Mg3, Mg7 was characterized by upregulation of DAM genes (*Spp1*, *Apoe*, *Cst7*, *Igf1*, *Lgals3bp*, *Apoc1*, *Lpl, Gpnmb, Itgax*)^31^ and by expression of genes observed in the response of microglia to neurodegeneration (*Gpnmb*, *Axl*, *Itgax*, *Spp1*, *Apoe*)^36^. Mg6 showed upregulation of immune genes (*Il1b*, *Nfkbiz*, *Cd83*, *Ccl4*) and downregulation of *Spp1* and *Lpl,* which suggests a pro-inflammatory phenotype, previously described in ischemic stroke and subarachnoid hemorrhage ^29,37^. In addition, Mg6 showed enriched gene expression signature of disease inflammatory macrophages (DIM; *Il1b*, *Cd83*, *Nfkbiz*, *Atf3*, *Ccl4*, *Egr1*, *Fosb*), indicating Mg6 cluster as DIM-like microglial cells (Fig. S5B)^38^. Pseudo-time trajectory predicted the progression of Sham microglia (Mg1) to either D02 or D14 microglia, whereas we could not observe a trajectory from D02 to D14 microglia (Fig. 2E).

**Figure 2.**
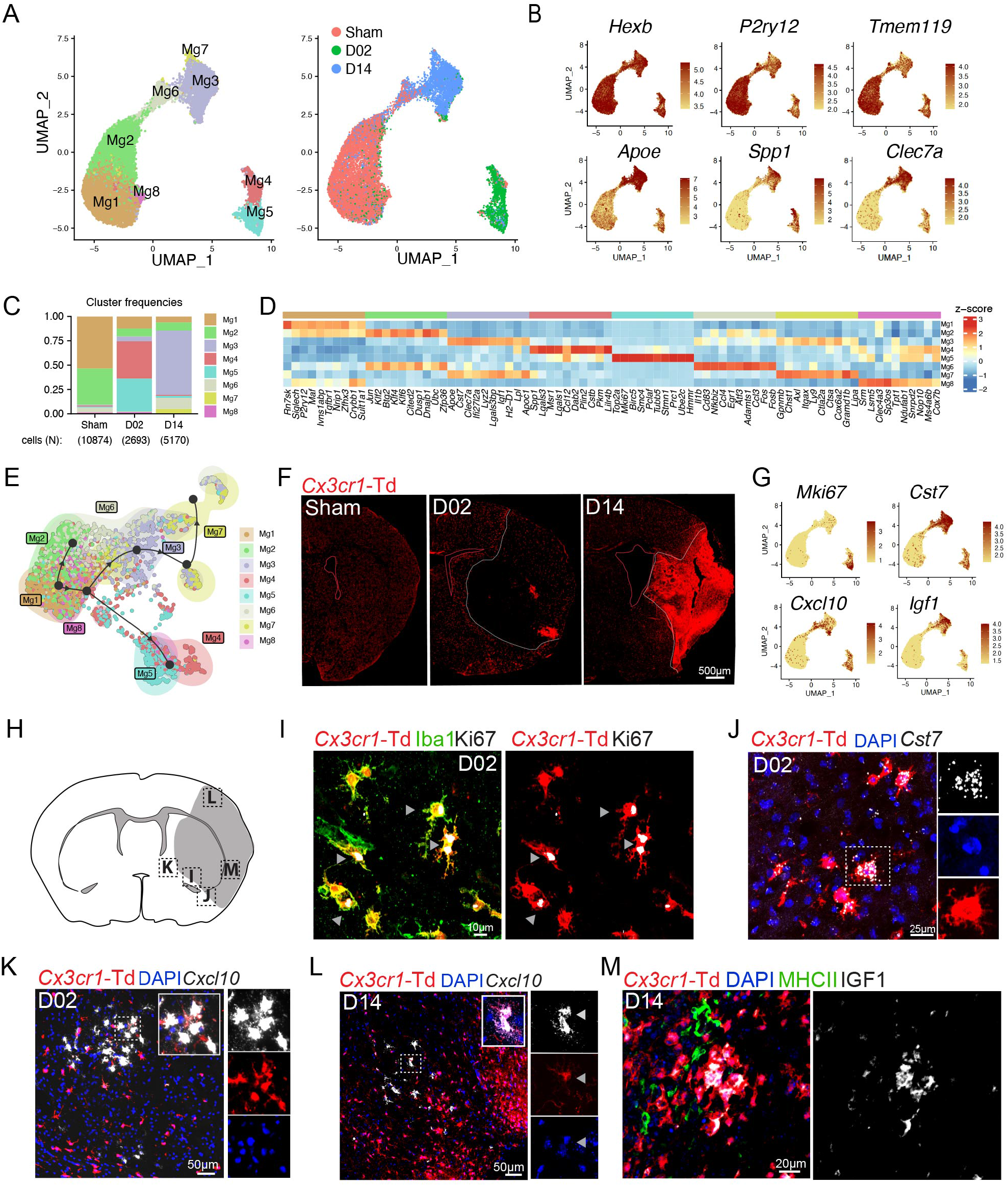
Subclustering analysis of microglia reveals altered transcriptional states through the acute and subacute stroke phases. **(A)** *Left:* UMAP analysis of merged Sham, D02 and D14 microglial single cell transcriptomes reveals 8 subclusters; *Right*: UMAP of overlaid time points reveals high segregation of microglial clusters among Sham, D02 and D14 groups. **(B)** Feature plots depicting single-cell gene expression of individual genes characterizing homeostatic and activated microglia. Scale bar represents log of normalized gene expression. **(C)** Bar graph showing relative frequencies of each microglial subcluster across Sham, D02 and D14 groups. **(D)** Heatmap displaying expression of the top 10 upregulated genes in each microglial cluster. Scale bar represents Z-score of average gene expression (log). **(D)** Heatmap displaying expression of the top 10 upregulated genes in each microglial cluster. Scale bar represents Z-score of average gene expression (log). **(E)** Pseudotime analysis of microglial subclusters showing transitional trajectories from Sham-Mg1 cluster to either Sham-Mg2, or D02 Mg4-5 clusters or D14 Mg3-7 clusters. **(F)** Representative fluorescence images of the cerebral hemisphere after Sham surgery or after 2 (D02) or 14 (D14) days MCA occlusion in *Cx3cr1*^CreERT2^:R26tdTomato mice. Microglia were identified as Td^+^ cells (red). Images show that the number of microglia decreased in the center of the lesion (white dashed outline) at D02 whereas its number increased at D14. **(G)** UMAP plots depicting single-cell gene expression of individual marker genes that characterize ischemic microglial clusters. Scale bar represents log of normalized gene expression. **(H)** Graphical representation of brain coronal section indicating anatomical regions where images I-L were acquired. **(I)** Immunofluorescence images showing Ki67 (white) expression by *Cx3cr1*-Td^+^(red) Iba1^+^(green) microglia 2 days after MCA occlusion. Arrowheads indicate Ki67 staining. **(J)** RNAscope in situ hybridization fluorescence (FISH) and immunofluorescence (IF) images validating *Cst7* (white) expression in D02 *Cx3cr1*-Td^+^ microglia (Td^+^, red). Nuclei are stained with DAPI (blue). **(K, L)** FISH images showing *Cxcl10* (white) expression in D02 (K) and D14 (L) *Cx3cr1*-Td^+^ microglia (Td^+^, red). Nuclei are stained with DAPI (blue). **(M)** IF images showing IGF1 (white) expression by *Cx3cr1*-Td^+^(red) MHCII^-^ (green) microglia 14 days after MCA occlusion.

Microglia undergo significant morphological and functional differentiation after cerebral ischemia, which could confound discrimination from infiltrating MdC ^39,40^. To verify microglial identity we used tamoxifen-inducible *Cx3cr1*^CreERT2^:R26tdTomato (*Cx3cr1*-Td) mice, a model that distinguishes Td^+^ microglia and BAM from hematopoietic myeloid cells ^41-43^. We subjected *Cx3cr1*-Td mice that had been injected with tamoxifen 6-8 weeks prior to Sham, D02 or D14 stroke and performed scRNA-seq on Td^+^ brain cells sorted by flow cytometry. Cells were annotated using the WT brain dataset as a reference. Clustering analysis of *tdTomato* expressing cells revealed that microglia constituted the majority of Td^+^ cells (92.7%) followed by BAM (2.9%), MdC (2.0%) and DC (1.3%). Considering that the number of MdC is nine times higher than the number of BAM, and that the number of DC is three times higher, we infer that the majority of BAM were Td^+^ while only a small proportion of MdC and DC retained Td expression (Fig. S5C).

To investigate microglia spatial distribution, we examined brains of *Cx3cr1*-Td stroke mice by histology and used FISH-IF to spatially map major microglia clusters (Fig. 2F, H-M). We observed that *Cx3cr1*-Td^+^ cells were significantly reduced in the ischemic core at D02, whereas they accumulated in the core at D14 (Fig. 2F). We observed colocalization of *Cx3cr1*-Td^+^Iba1^+^ microglia with the mitotic marker Ki67 at the infarct border at D02 (Fig. 2I; Fig. S6A), suggesting that proliferative microglia contribute to the repopulation of the ischemic territory seen at D14.

We found the DAM marker *Cst7* ^31,44^ in both D02 and D14 microglia. *Cst7* colocalized with ramified microglial cells at the infarct border and with ameboid cells in the ischemic core (Fig. 2J; Fig. S6B) suggesting that cell morphology was not associated with a DAM signature. We also identified *Cxcl10* and *Igf1* among the DEGs that were upregulated in microglia after ischemia (Fig. 2G). Microglial induction of the inflammatory chemokine *Cxcl10,* a IFN type I stimulated gene, has previously been reported in models of traumatic and ischemic brain injury, EAE and AD ^29,45-48^, while IGF-1 expressing microglia has been reported to be associated with a pro-neurogenic phenotype after stroke in rats ^49^. We observed that *Cxcl10* was upregulated at D02 and D14, whereas *Igf1* was mainly induced at D14 (Fig. 2G). By histology, we also found *Cxcl10* expressing microglia in the infarct border at D02 and in the lesioned tissue at D14, while IGF1^+^ microglia were found in the ischemic core (Fig. K-M; Fig. S6C-D). In some instances, Td^+^*Cxcl10*^+^ microglia were found near Td^-^*Cxcl10^+^* cells, suggesting microglia and non-microglial cells expressing *Cxcl10* organize into discrete cell clusters (Fig. 2K).

### Border-associated macrophages exhibit distinct transcriptional states early after stroke

We identified five BAM clusters (Fig. 3A). BAM1 and BAM2 were the most abundant clusters in all groups (Fig. 3C). BAM1 showed transcriptional signature of subdural BAM (*Cd209f, Ccl24*, *Clec10a*, *Slc40a1*, *Stab1*) ^50^. Some of these genes have been found in homeostatic BAM and are related to leukocyte recruitment (*Ccl24*), phagocytosis (*Cd209f*) and iron metabolism (*Slc40a1, Cp*) ^51,52^. BAM2 showed the highest separation from all other BAM clusters and displayed upregulation of canonical marker genes of choroid plexus macrophages (ChMp) such MHC class II genes (*H2-Ab1*, *H2-Eb1*, *H2-Aa*, *H2-DMb1*, *H2-DMa*), the MHCII-associated gene *Cd74*, *Ccr2*, and low level of the perivascular macrophages (PVM) marker *Lyve1* (Fig. 3B,D) ^52-55^. BAM3, which was less frequent at D14, featured high expression of genes coding for enzymes *Wwp1*, *Abhd12*, *Dhrs3, Hpgd,* and transferrin (*Trf*). BAM4 was largely confined to D02 and was characterized by the upregulation of genes linked to macrophage/microglia activation in either ischemic stroke or other neurodegenerative diseases (*Spp1*, *Ccl8*, *Cstb, Lgals1*) ^50,56,57^. BAM5 was associated with induction of genes related to lipid metabolism (*Fabp5, Lgals3)* ^58^ and damage-associated molecular patterns (DAMPs)-recognition molecules (*S100A4, Lgals3, Clec4d, Clec4e*) ^59^.

**Figure 3.**
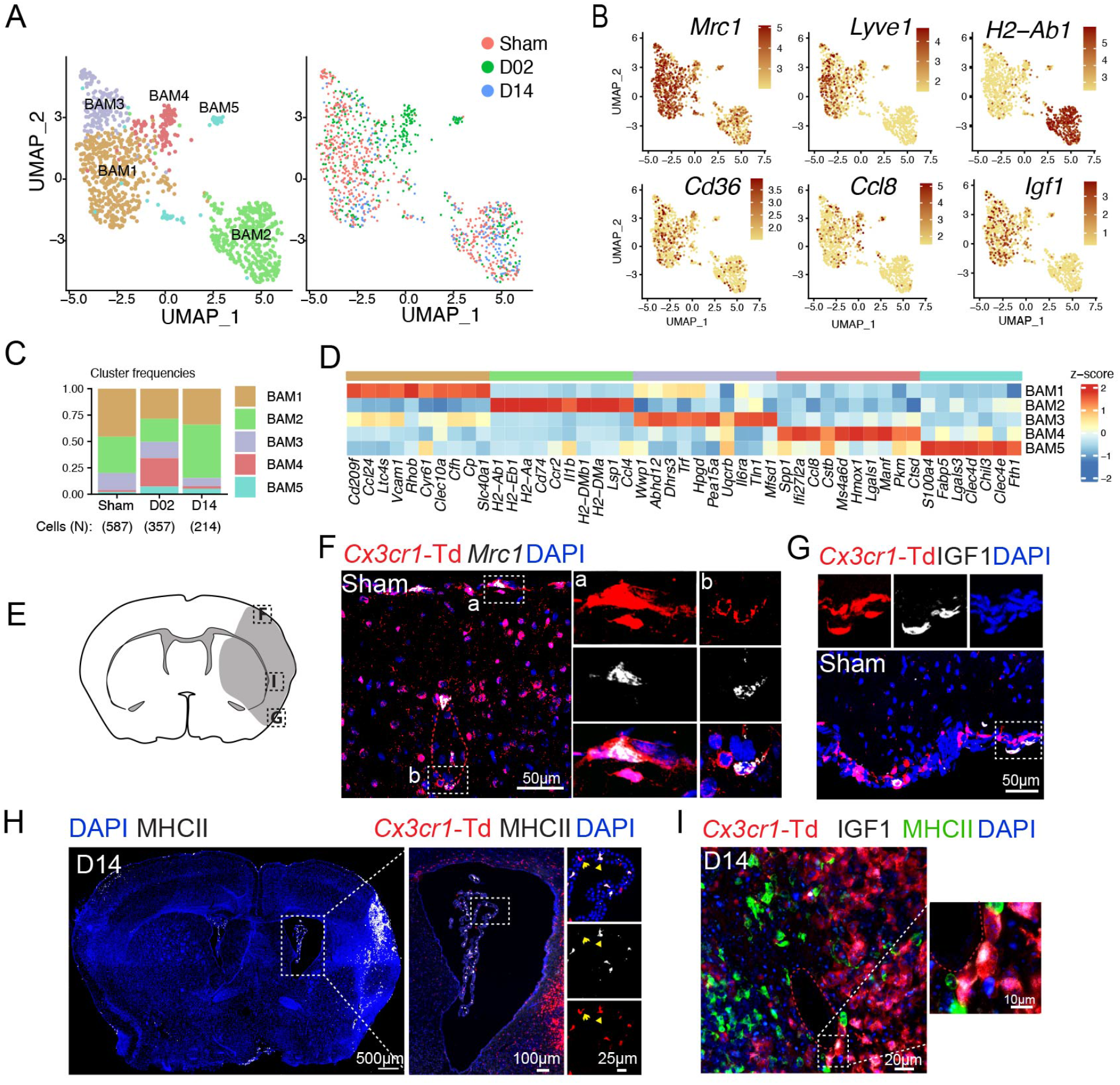
Transcriptional changes of border-associated macrophages (BAM) after stroke. **(A)** *Left*: UMAP analysis of merged Sham, D02 and D14 BAM transcriptomes discloses 5 subclusters; *Right*: UMAP of overlaid time points shows overlap of clusters among Sham, D02 and D14 groups, except for BAM4, which is confined to D02. **(B)** UMAP plots depicting expression of individual marker genes for BAM (*Mrc1*, *Cd36*), meningeal and perivascular BAM (*Lyve1*), choroid plexus BAM (*H2-Ab1*) and activated meningeal and perivascular BAM (*Ccl8*, *Igf1*). Scale bar represents log of normalized gene expression. **(C)** Bar graph showing relative frequencies of BAM subclusters across Sham, D02 and D14 groups. **(D)** Heatmap displaying expression of the top 10 upregulated genes in each BAM cluster. Scale bar represents Z-score of average gene expression (log). **(E)** Graphical representation of brain coronal section indicating anatomical regions where images F-I were acquired. **(F)** RNAscope in situ hybridization fluorescence (FISH) combined with immunofluorescence (IF) images of brain cortical areas showing *Mrc1* expression (white) in resident macrophages (Td^+^, red) on the brain surface and around vessels of *Cx3cr1*-Td^+^ mice 2 days after Sham surgery (Sham). Nuclei are stained with DAPI (blue). **(G, I)** IF images validating IGF1 (white) expression by pial BAM (Td^+^, red) (**G**) in Sham *Cx3cr1*-Td+mice and in perivascular macrophages (PVM) (I) of the ischemic brain at D14 (*Middle and left*). **(H)** *Left*: Representative IF image of a whole brain section from a *Cx3cr1*-Td^+^ mouse subjected to 14 days of MCA occlusion (D14), showing MHCII^+^ cells (white, binary mask) localization and nuclear DAPI staining (blue); *Right*: IF images of magnified areas of the choroid plexus (ChP) showing MHCII expression (white) by ChP macrophages (Td^+^, red).

Utilizing *Cx3cr1*-Td mice, we mapped BAM on the surface of the brain or lining parenchymal blood vessels by *Mrc1* (coding for CD206) and Td co-detection. Consistent with the scRNA-seq data, MHC class II antigen was only detected in ChMp, while PVM and pial macrophages (pMp), but not ChMp, expressed IGF1 (Fig. 3B, E-I).

### Brain monocyte-derived cells arise from circulating inflammatory monocytes and diversify over time in the brain after stroke

After sub-setting and re-clustering, we identified 5 clusters of blood monocytes (Mo1-5). All the clusters except for Mo2, which was almost absent at D14, were present at all studied time points (Fig. 4A). Based on *Ly6c2* and *Cd36* expression, Mo clusters were annotated as inflammatory (*Ly6c2* high*, Cd36* low) or patrolling (*Ly6c2* low*, Cd36* high) monocytes (Geissmann et al., 2003; Marcovecchio et al., 2017). Inflammatory monocytes separated into 4 clusters (Mo1,2,4 and 5), whereas patrolling monocytes constituted a single cluster (Mo3) (Fig. 4B). Mo1 was most abundant at D14 and was characterized by increased expression of S100 proteins and chemokines (*S100a10*, *S100a4*, *Ccl9*, *Ccl6*). On the other hand, Mo2 was highest at D02 and showed transcriptional profile of ‘neutrophil-like’ Ly6C^hi^ monocytes (*Saa3*, *Mmp8, Lcn2, Wfdc21, Lrg1, Chil3*) ^60^. In addition to *Cd36,* Mo3 expressed several genes characteristic of patrolling monocytes (*Ear2*, *Treml4*, *Eno3*, *Aceas*) ^60^.

**Figure 4.**
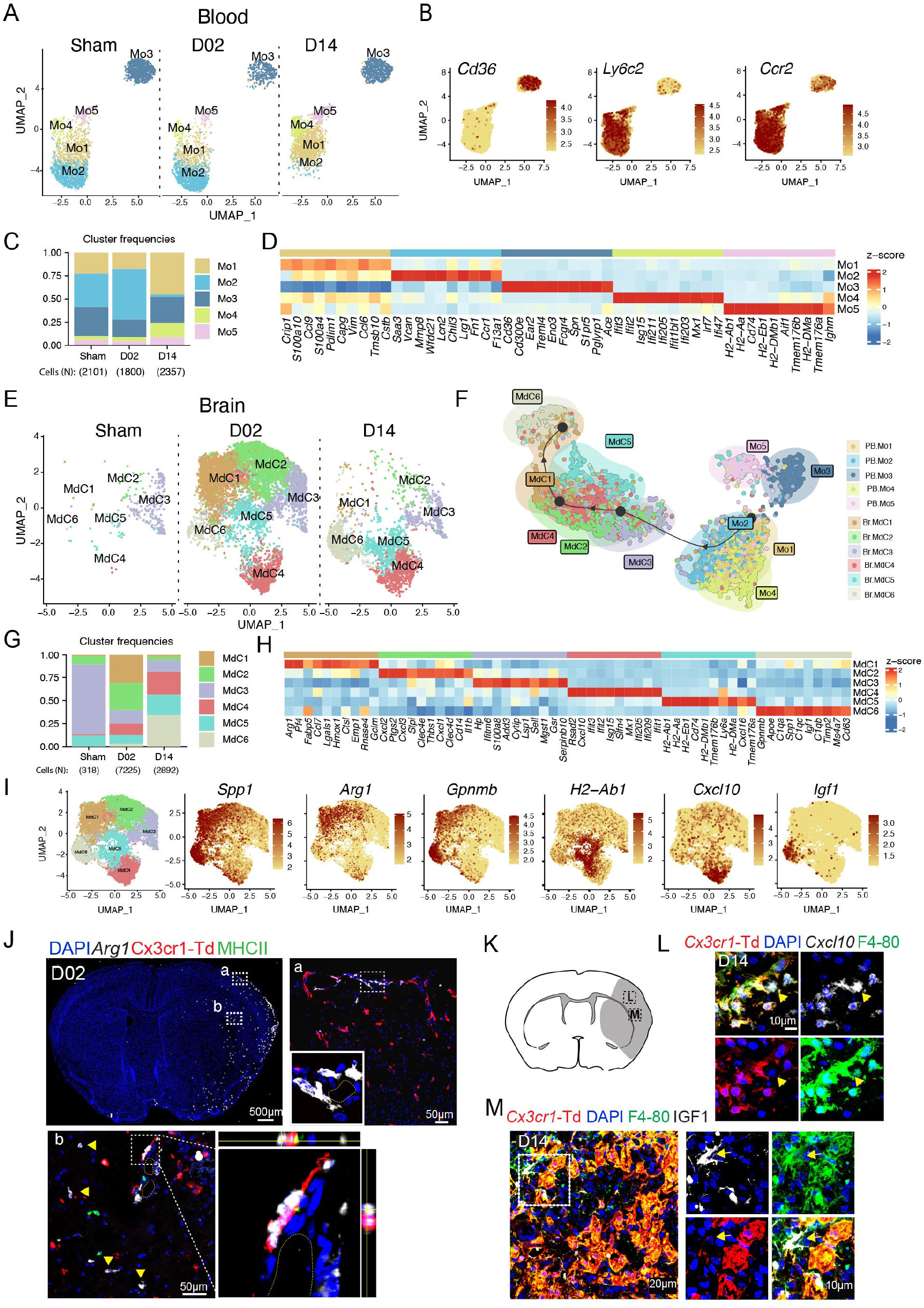
Inflammatory blood monocytes give rise to infiltrating brain macrophages (MdC) after stroke. **(A)** UMAP plot of peripheral blood monocyte transcriptomes (Mo) in control surgery (Sham) or stroke mice 2 or 14 days (D02, D14) after injury, showing 5 monocyte clusters (Mo1-5). **(B)** UMAP plots depicting single-cell gene expression of individual marker genes for monocytes (*Ccr2*) and for inflammatory vs. patrolling monocyte identification (*Cd36*, *Ly6c2*). Scale bar represents log of normalized gene expression. **(C)** Bar graph showing relative frequencies of monocytes subclusters across Sham, D02 and D14 groups. **(D)** Heatmap displaying expression of the top 10 upregulated genes in each monocyte cluster. Scale bar represents Z-score of average gene expression (log). **(E)** UMAP plot of brain monocyte derived cell (MdC) transcriptomes for each studied time point shows 6 clusters. **(F)** Slingshot trajectory analysis of peripheral blood monocytes (Mo) and brain macrophages (Mdc) subclusters. Each point is a cell and is colored according to its cluster identity shown in (**A**) and (**E**). The inferred trajectory shows transition from blood inflammatory monocyte clusters Mo1 and Mo2 to fully differentiated brain macrophages MdC1 and MdC6. **(G)** Bar graph showing relative frequencies of MdC subclusters across Sham, D02 and D14 groups. **(H)** Heatmap displaying expression of the top 10 upregulated genes in each MdC subcluster. Scale bar represents Z-score of average gene expression (log). **(I)** UMAP plots displaying expression of marker genes for each identified MdC brain clusters. Scale bar represents Z-score of average gene expression (log). **(J)** RNAscope fluorescent in situ hybridization (FISH) for *Arg1* (white) combined with immunofluorescence (IF) brain images from *Cx3cr1*-Td^+^ mice 2 days after stroke. *Top left*: Overview of *Arg1* expression (binary mask) in a whole brain section co-stained for DAPI (blue), showing high upregulation of *Arg1* in cortical areas of the ischemic hemisphere. *Top right and bottom*: images of magnified areas (a) and (b) showing localization of *Arg1* in *Cx3cr1*-Td^-^ and MHCII^-^ cells. *Bottom right*: orthogonal view of magnified area showing *Arg1* expression in an adjacent cell to a PVM (Td^+^, red). **(K)** Graphical representation of brain coronal section indicating anatomical regions where L and M images were acquired. **(L)** FISH-IF images of the ischemic cortical region in D14*-Cx3cr1*-Td^+^ mice showing *Cxcl10* expression (white) by MdC (Td^-^, F4-80^+^ cells, yellow arrow) near microglia (Td^+^, red; F4/80^+^, green) expressing *Cxcl10.* Nuclei are stained with DAPI (blue). **(M)** IF images of the ischemic cortical region in D14*-Cx3cr1*-Td^+^ mice validating IGF1 expression (white) by MdC. *Left*: Image shows IGF1 expression by MdC (Td^-^, F4/80^+^, green) on brain surface. *Right*: Image shows clusters of mixed microglia (Td^+^, red) and MdC (Td^-^, F4/80^+^, green) expressing IGF1.

Furthermore, Mo4 was characterized by an interferon-stimulated gene (ISG) signature (*Ifit3*, *Ifit2*, *Isg15*, *Ifi211*, *Ifi205*, *Ifit1bl1*, *Ifi203*) and Mo5 by high expression of MHC class II genes (*H2-Ab1*, *H2-Eb1*, *H2-Aa*, *H2-DMb1*, *H2-DMa*) and *Cd74,* resembling monocyte-derived dendritic cells ^61^ (Fig. 4C-D).

Sub-clustering of brain MdC identified six populations. Sham brain displayed low numbers of MdC and most of them (∼75%) clustered as MdC3, a cluster closely related to blood monocytes (*Serpinb10, Plac8, Sell*) ^62^(Fig. 4E and 4G, Fig. S7A-B). MdC1 and MdC2 were the predominant clusters at D02 and MdC6 was the major cluster at D14. MdC1 expressed a gene signature (*Fabp5, Spp1, Gpnmb, Ctsl, Cd63, Ctsb, Ctsd, Arg1*) that has been found in stroke-associated macrophages ^16^ and which is reminiscent of the gene signature found in either foamy macrophages in atherosclerotic plaques or lipid-associated macrophages in myocardial infarct (Fig. 4H, Fig. S7B) ^63,64^. MdC2 showed upregulation of neutrophil chemoattractants (*Cxcl1, Cxcl2, Cxcl3)* and pro-inflammatory genes (*Ptgs2, Il1b, Clec4e*), suggesting a role in driving post-stroke inflammation. Like MdC1, MdC6 showed increased expression of genes characteristic of stroke-associated macrophages and foamy macrophages (*Gpnmb, Spp1, Cd63, Trem2* and *Fabp5*), but differ from MdC1 in that they were enriched for the repair growth factor *Igf1* ^65^. MdC4 cells showed an ISG signature (*Rsad2, Cxcl10*, *Ifi205*, *Ifit2*, *Ifit1*, *Ifit3*, *Ifit1bl1*, *Isg15*) (Fig. S7B), while MdC5 exhibited upregulation of MHC class II associated genes (*H2-Ab1, H2-Aa, H2-Eb1, Cd74, H2-DMb1*), a signature which identifies this cluster as monocyte derived-DC ^61^.

Slingshot trajectory analysis showed sequential transition of inflammatory peripheral blood Mo1 and Mo2 clusters to brain MdC3 cluster, followed by MdC2, MdC4 and MdC1 transition and final differentiation into MdC6. MdC5 appeared as the only MdC cluster dissociated from the trajectory (Fig. 4F). Neither patrolling Mo3 nor Mo5 were identified as MdC sources. Spearman correlation analysis between Mo and MdC clusters also showed proximity of blood monocytes to MdC3 and, consistent with the trajectory analysis, patrolling Mo3 cluster showed the lowest correlation with all brain MdC clusters (Fig. S7A). Thus, these analyses suggest that MdC in the inflamed brain derive from inflammatory monocytes, as recently described ^60^ and that both MdC1 and MdC6 constituted bona-fide tissue macrophages, while MdC2, MdC3, and MdC4 were clusters associated with transitional phenotypes.

*Spp1* (osteopontin) and *Gpnmb* (osteoactivin) were preferentially upregulated in fully differentiated MdC1 and MdC6 clusters (Fig. 4H-I). Upregulation of these two genes in brain macrophages has been linked to tissue regeneration and neuroprotection after stroke ^66,67^. MdC1 was characterized by *Arg1* expression, a gene associated with efferocytosis ^68^. Detection of *Arg1* by combined FISH-IF revealed *Arg1* increased at D02 in infiltrating macrophages (*Cx3cr1*-Td^-^ cells) located on the brain surface or structures resembling parenchymal blood vessel as previously reported ^69^ (Fig4. J). In addition, we found upregulation of *Cxcl10* at both D02 and D14 (MdC4) and of *Igf1* at D14 (MdC6), similar to the one observed in microglia (Fig. 4K-M, Fig. S7C-D). Many of the *Cxcl10* expressing MdC were organized in cell clusters together with *Cxcl10* producing microglia indicating that local environmental cues in selected brain regions might drive the ISG phenotype in MdC and microglia.

### Endothelial cells subclusters display a reactive signature early after stroke identified by *Lrg1* expression

EC segregated into nine subclusters (EC1-9), which by marker gene expression could be attributed to four arteriovenous segments: EC1 (venous capillaries; *Car4, Tfrc*), EC6 (large veins; *Vcam1*, *Cfh*, *Scl38a5*), EC3 (arterial capillaries; *Fos, Fosb*), EC4 (arteries; *Gkn3, Mgp*, *Stmn2*, *Bmx*), and EC7 (arteries; *Clu*, *Cdh13, Mgp, Stmn2, Bmx*) ^70-72^. EC2, EC6 and EC9, which clustered together in the UMAP space, showed upregulation of the endothelial venule marker *Lrg1,* a gene associated with angiogenesis after ischemic stroke ^73^. In addition, EC6 was defined by high expression of the venular atypical chemokine receptor *Ackr1* ^74^, lipocalin-2 (*Lcn2),* which has been implicated in stroke damage ^75^, and, consistent with a venous phenotype, adhesion molecules *Icam1* and *Vcam1*. Attesting to their reactive state, EC2, which was the predominant cluster at D02, was defined by expression of *Ecscr*, a gene related to EC migration and vessel formation ^76^*, Anxa2* and *S100a11* implicated in endothelial fibrinolysis ^77,78^, and *Scgb3a1*, a endothelial secreted protein crucial for metastasis ^79^. EC9 was characterized by genes involved in EC metabolic reprogramming (*Tyms, Dut, Dctpp1*) ^80^ and proliferation (*Pimreg, Pclaf*). EC8 showed signature of fenestrated brain vascular ECs, likely stemming from choroid plexus EC (*Plvap, Plpp3, Igfbp3, Plpp1, Cd24a, Ldb2*) ^81^. Moreover, EC5 displayed increased ISG gene expression (*Ifit3*, *Isg15*, *Rsad2*, *Ifit1*, *Usp18*, *Irf7*, *Ifit3b*, *Iigp1*, *Ifi44*, *Gbp2*, *Ifit2*, *Gbp3*) (Fig. 5A-D). It has been reported that brain EC express IFN type I inducible genes at homeostasis, and that after stroke, EC show reduced ISG expression ^82^. Despite our data showing significant ISG scores in the Sham group, they were further increased at 2 or 14 days after stroke (Fig. S8). We also found that ECs, particularly EC1, EC3 and EC5, exhibited high *Igf1r* expression (Fig. 5B). *Igf1r* levels are elevated in brain endothelial cells compared to peripheral tissues ^83^. Because we found *Igf1* expression in microglia, MdC, and BAM, we performed CellChat analysis to investigate interactions between ECs and IGF1 producing cells. Whereas *Igf1-Igf1r* communication occurred between BAM and EC in Sham and D02 groups, both microglia and MdC6 contributed to *Igf1-Igf1r* signaling network at D14 (Fig. 5E).

**Figure 5.**
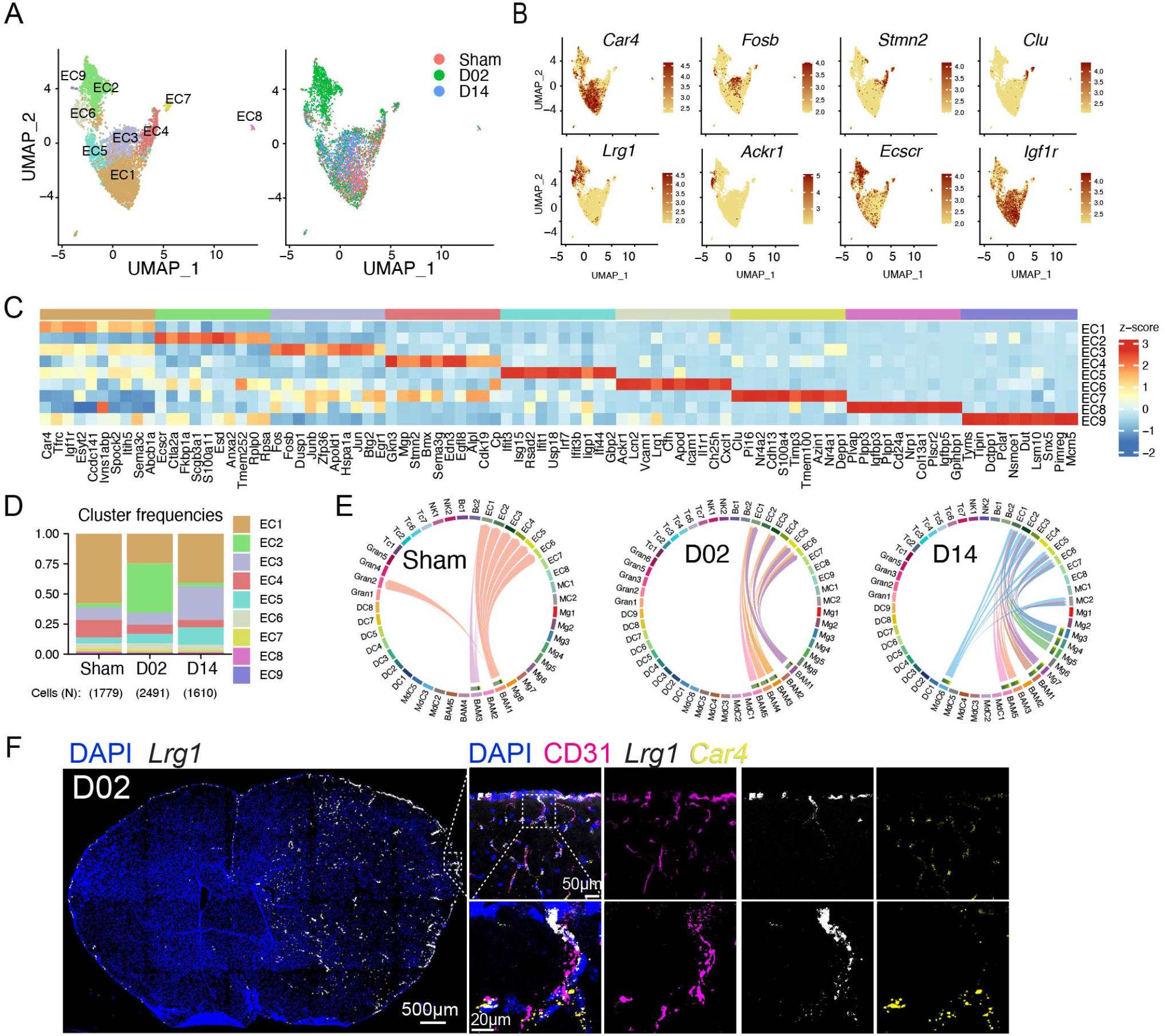
Endothelial cell (ECs) transcriptional changes and *Igf1r* signaling after stroke. **(A)** *Left:* UMAP analysis of merged Sham, D02 and D14 EC transcriptomes discloses 9 subclusters; *Right*: UMAP of overlaid time points shows dissociation of EC2 cluster at D02 and general overlap of all the other clusters across Sham, D02 and D14 groups. **(B)** UMAP plots depicting single-cell gene expression of individual marker genes for capillary (*Car4*, *Fosb*), arterial (*Stmn2, Clu)* and venular (Lrg1, *Ackr1*) ECs and for the endothelial receptors *Ecscr* and *Igf1r*. Scale bar represents log of normalized gene expression. **(C)** Heatmap displaying scaled differential expression of the top 10 upregulated genes in each EC cluster. Scale bar represents Z-score of average gene expression (log). **(D)** Bar graph showing relative frequencies of EC subclusters across Sham, D02 and D14 groups. **(E)** Chord plots showing CellChat inferred ligand-receptor interactions between *Igfr1* and *Igf1* in Sham, D02 and D14 stroke mice. The strength of the interaction is indicated by the edge thickness. The color of the chord matches the cell cluster color sending the signal (Igf1). The number of cell cluster receptors (*Igfr1*), and their weight in the interactions, is indicated by the color-matched stacked bar next to each sender cluster. **(F)** RNAscope in situ hybridization fluorescence (FISH) combined with immunofluorescence (IF) of *Lrg1*. *Left*: Overview of *Lrg1* expression (binary mask, white) in a whole brain section co-stained for DAPI (blue), showing high upregulation of *Lrg1* in the ischemic hemisphere *Right*: magnified images of brain cortical areas showing *Lrg1* (white) colocalization with the EC marker CD31 (magenta) but not with the capillary marker *Car4* (yellow). Nuclei are stained with DAPI (blue).

FISH-IF analysis for the venule marker *Lrg1* ^70^ revealed strong upregulation of vascular *Lrg1* in the ischemic hemisphere (Fig. 5F). Co-detection of *Lrg1* and *Car4*, a marker for venule capillaries, showed low overlap between both markers, indicating that *Lrg1* was mainly induced in larger veins. LRG1, is a secreted glycoprotein primarily synthesized by neutrophils and hepatocytes, however, in conditions of altered homeostasis its transcription is upregulated in endothelial cells ^84^. Indeed, LRG1 expression increased in the mouse brain 1-3 days after MCA occlusion ^85^, although its localization in the endothelium was not demonstrated. Among its described vascular effects, LRG1 has been linked to pathological angiogenesis through modulation of endothelial cell TGFβ signaling ^86^ and it has also been described as a predictor for arterial stiffness and endothelial dysfunction ^84,87^. On the other hand, LRG1, produced from myeloid cells, contributed to cardiac repair after myocardial infarction in mice ^88^. Although we did not explore LRG1 activity in our stroke model, our data identify *Lrg1* as a powerful marker of a specific subset of venular brain EC that respond early after ischemia.

### Transcriptional changes of brain granulocytes indicate continuous recruitment from the circulation

We performed subclustering and cell trajectory analyses to investigate whether granulocytes states evolve following a transcriptional continuum defined by either their migration from periphery to the injured brain or by their transition over the ischemic-reperfusion time. We detected 6 clusters of blood neutrophils, which relative frequencies were similar between Sham and D02 groups whereas at D14, blood Neu1, 2 and 4 clusters were practically absent and Neu3 became the predominant cluster (Fig. 6A-C). Neu1 showed increased expression of *Cd14*, *Cebpb, Il1b* and *Lrg1.* Neu2 was characterized by upregulation of genes described in immature neutrophils (*Mmp8*, *Retnlg*, *Lcn2*, *Ly6g*) ^89^, and Neu4 cluster was enriched in stefinA family genes (*Stfa1, Stfa2, Stfa3, Stfa2l1, BC100530*), which are cytoplasmic inhibitors of proteases such as cathepsins ^90^. On the other hand, Neu3, the predominant cluster at D14, was characterized by genes associated with neutrophil infiltration (*Ninj1, Cd300c2)* ^91^, cell growth inhibition (*Lst1*, *Creg1*) or defining a mature neutrophil state (*Cd101*) ^92^.

**Figure 6.**
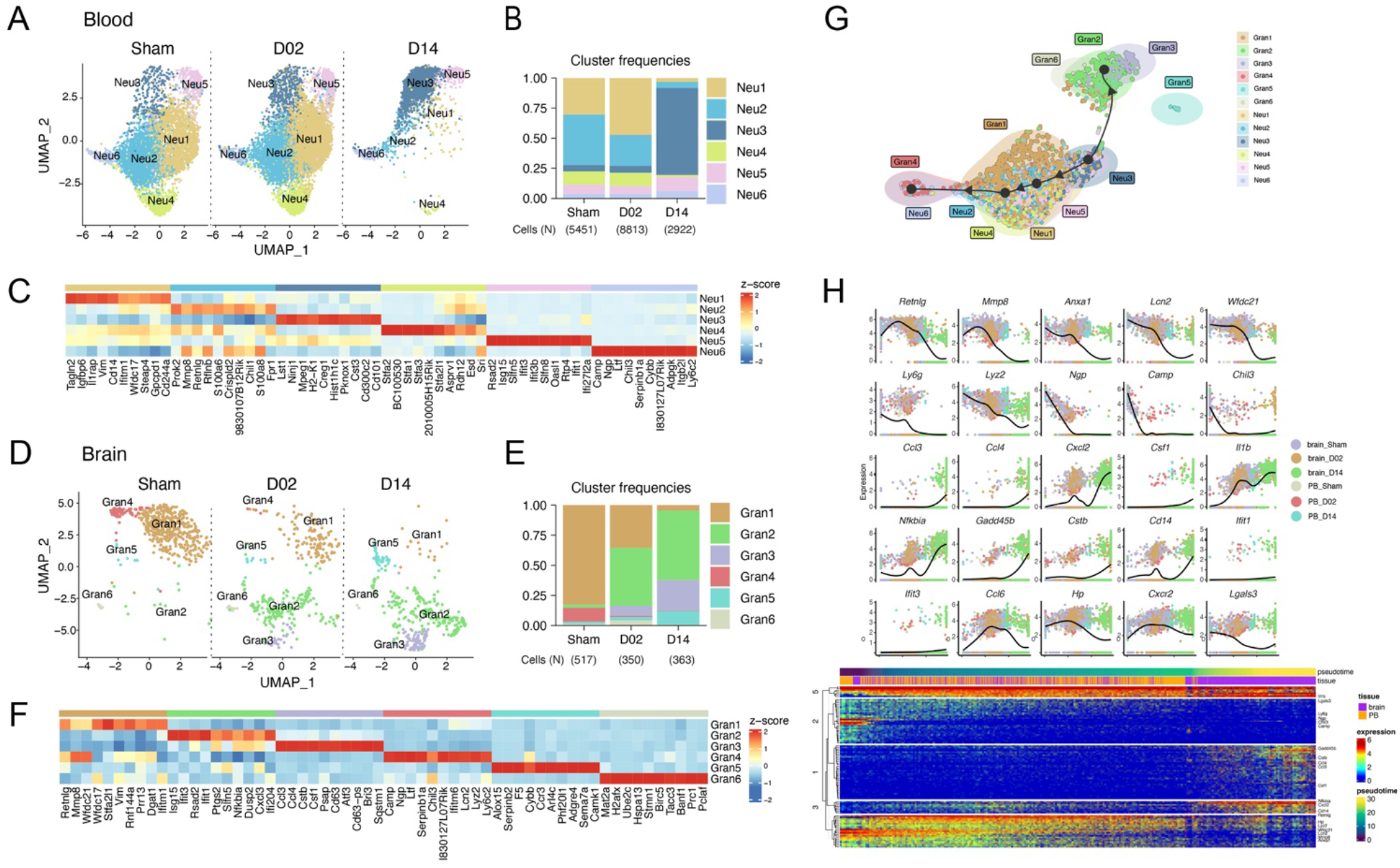
Granulocyte transcriptional changes through ischemia-reperfusion. **(A, D)** UMAP plots of peripheral blood neutrophil (Neu) **(A)** and brain granulocyte (Gran)**(C)** transcriptomes by studied time point: control surgery or stroke mice 2 or 14 days after injury (Sham, D02, D14). Six clusters were identified in each data set (Neu1-6; Gran1-6). **(B, E)** Bar graph showing relative frequencies of peripheral blood Neu **(B)** and brain Gran **(D)** clusters across Sham, D02 and D14 groups. **(C, F)** Heatmap displaying scaled differential expression of the top 10 upregulated genes in each peripheral blood Neu **(E)** and brain Gran **(F)** clusters. Scale bar represents Z-score of average gene expression (log). **(G)** Slingshot trajectory of combined peripheral blood neutrophils (Neu) and brain granulocytes (Gran) showing predicted cluster transition. Each point is a cell and is colored according to its cluster identity. The analyses point out transcriptional states changes of blood and brain granulocyte through the ischemic-reperfusion time. **(H)** *Top*: Expression of single gene plotted as a function of pseudotime. Dot plots show expression levels of top cluster genes of combined blood Neu and brain Gran data sets along ischemic-reperfusion pseudotime. Each dot represents the expression levels (log) for each gene in a cell and is colored according to the group. Lines show average expression. *Bottom*: The top 100 genes that specifically covary with pseudotime were identified using generalized additive models and the log normalized expression values were plotted along the pseudotime axis. The location of genes plotted above is indicated.

Regarding the brain, granulocyte subclustering also identified 6 subsets. Most of the sham brain granulocytes fell into Gran1 and 4 clusters, which, similar to blood Neu2, were characterized by the upregulation of the immature neutrophil marker genes *Retnlg, Mmp8,* and *Wfdc21* ^89^. In addition, Gran4 was specifically characterized by induction of genes associated with early stage neutrophil development (*Camp, Ltf, Chil3, Lcn2*) ^89^. We observed that Gran1 and 4 frequencies gradually decreased over the ischemia-reperfusion time, whereas the frequencies of Gran2, which showed upregulation of ISG genes (*Isg15*, *Ifit3*, *Rsad2*, *Ifit*, *Ifi204*) and Gran 3, characterized by the cytokine genes (*Ccl3*, *Ccl4*, *Csf1*) augmented progressively over D02 and D14. Furthermore, we detected two small clusters, Gran5, which also contained eosionophil markers (*Ccr3*, *Alox15*), and Gran6, which showed upregulation of the cell cycle genes *Pclaf*, *Banf1* and *Prc1* (Fig. 6D-F).

Slingshot trajectory analysis of combined blood (Neu) and brain (Gran) granulocytes, indicated similarities between Gran1, which is the major Sham brain granulocyte cluster, and Neu1, 2, and 4, which were the main blood clusters at Sham and D02. On the contrary, Gran2 and 3 were disconnected from sham brain granulocytes (Gran1 and 4) and closer to blood Neu3, the predominat blood granulocyte cluster at D14 (Fig. 6G). Thus, this analysis suggests that, in contrary to MdC, granulocyte transcriptomic states are not developing within the tissue but suggests recruitment of brain infiltrating granulocytes from the circulating pool at early and late phases after ischemia-reperfusion. To further investigate granulocyte transcriptomic patterns over the ischemia-reperfusion time, we plotted gene expression as a function of pseudotime of selected genes that showed the strongest correlations with granulocyte transcriptomic shifts (Fig. 6H). We found that the pseudotime trajectory was characterized by the early expression of *Retnlg*, *Mmp8*, *Ly6g, Anxa1 and Lcn2* genes, top marker genes describing Sham brain granulocytes and Sham and D02 blood neutrophils. Conversely, late expressed genes included *Ccl3, Ccl4, Csf1, Gadd45b* in D14 brain granulocytes, suggesting different functions of brain neutrophils under homeostatic conditions, the early and late phases of ischemic injury.

### Transcriptional changes in dendritic cells (DC)

We identified nine DC subclusters in the stroke mouse brain. Low numbers of DC were found in brains of Sham mice, which gradually increased over the ischemia-reperfusion time (Fig. 7A-B). Expression signatures of the clusters were related to the five main DC populations based on canonical markers: conventional cDC1 (*Xrc1*, *Clec9a*) and cDC2 (*CD209a, Sirpa* (CD172a)), the plasmacytoid DC (pDC; *Siglech*, *Ccr9*, *Bst2*), a subpopulation with high *Ccr7* expression representing migratory DC (migDC), and monocyte-derived DC (moDC; *Ms4a7, Lyz2*) ^20,55,93-97^ (Fig. 7C-E). We confirmed by flow cytometry the presence of DC (CD45^hi^F480^-^Lin-CD11c^+^MHCII^+^) subtypes cDC1 (XCR1^+^), cDC2 (CD172a^+^ and CD209a^+^) and migDC (CCR7^+^) in the ischemic brain (Fig. 7F). cDC1 were comprised by clusters DC2 and DC9. DC2 showed upregulation of *Irf8* transcription factor, which is required for the full development of cDC1^98,99^, whereas DC9 additionally showed transcriptional signature of cDC1 dividing cells (*Lig1*, *Top2a*, *Mki67*, *Pcna*) ^100^. cDC2, the largest population of brain DC, comprised DC1, DC3, DC6 and DC7 clusters. DC1, the cDC2 cluster displaying the highest frequency at D14, showed prominent upregularion of genes associated with antigen presentation (*Cd72*, *H2-Oa*, *H2-DMb2*), while DC7 showed expression of scavenger receptors (*Clec4b1, Mrc1, Cd209a*). DC4 exhibited high expression of monocyte/macrophage marker genes (*Lyz2*, *Csf1r,* Apoe, genes encoding complement C1q chains *C1qa*, *C1qb*, *C1qc, Ms4a7, Trem2 and Cd14)* identifying them as moDC ^101^ and DC6 was characterized by the upregulation of ISG genes (*Ifit3, Rsad2, Cxcl10, Ifit2, Ifi204, Ifit1bl1, Isg15, Usp18*). migDC were composed of a single subcluster (DC5) which, in addition to *Ccr7,* expressed other genes characteristic for migDC including *Fscn1*, *Tmem123*, *Ccl22,* and *Socs2* ^55,102^. DC8 classified as plasmacytoid DC based on the expression of *Ccr9*, *Bst1, Il3ra, Siglech, Irf7, Ly6d* ^103^ (Fig. 7E).

**Figure 7.**
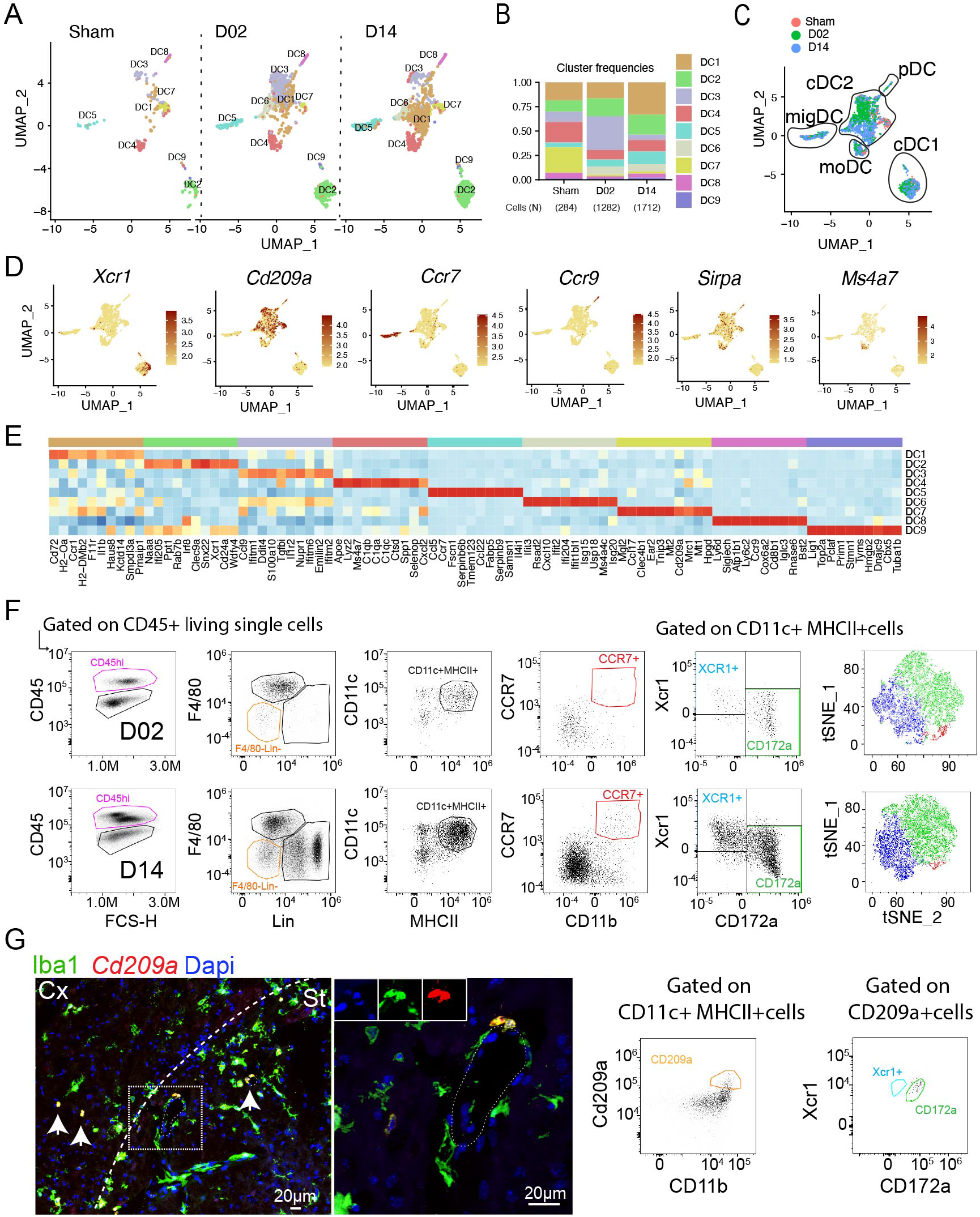
Transcriptional changes in brain dendritic cells. **(A)** UMAP plots of brain dendritic cells (DC) transcriptomes for each studied time point identifies 9 clusters (DC1-9). **(B)** Bar graph showing relative frequencies of DC subclusters across Sham, D02 and D14 groups. **(C)** UMAP of 3 color-coded time point overlay of brain DC. Classification of the clusters in DC subtypes is indicated by diagrams: conventional DC (cDC1, cDC2), migratory (mDC), monocyte derived-DC (moDC) and plasmacytoid DC (pDC). **(D)** UMAP plots displaying expression of marker genes for each identified DC cluster in the brain. Scale bar represents log of normalized gene expression. **(E)** Heatmap displaying scaled differential expression of the top 10 upregulated genes in each DC subcluster. Scale bar represents Z-score of average gene expression (log). **(F)** Flow cytometry analysis validating brain cDC1 (XCR1+), cDC2 (CD172a+) and migDC (CCR7+) subtypes identified by scRNA-seq after stroke. **(G)** *Left:* FISH of *Cd209a* (red) expression in the brain, combined with IF for Iba1 (green) and nuclear satining with DAPI, showing *Cd209a^+^*Iba1^+^ cells around the vessels. Cx: cortex; St: striatum. *Right*: Flow cytometry analysis showing double positive CD209a^+^CD172a^+^ cells.

Regardig peripheral blood, we detected few DC as compared to other leukocyte types (Fig.1B). Subclustering analysis and automated cell type annotation identified 5 clusters which corresponded to cDC2 (DC1, DC4), moDC (DC2), and pDC (DC5) (Fig. S9). Whereas the number of blood cDC2 clusters DC1, DC2 and DC3 increased over the ischemic-reperfusion time, the number of DC5 (pDC) remained unchanged (Fig. S9A)

### Transcriptional changes in lymphoid cells

Brain-associated T cells split into seven clusters (Fig. 8A). T cells were more numerous at D14 as compared to Sham and D02 (Fig. 8B). Expression of *Cd3d* identified all but one cluster (Tc6) as bona fide T cells (Fig. 8E). Tc6 was classified as type 2 innate lymphoid cells (ILC2) based on the expression of *Il1rl1*, *Gata3*, *Areg*, and *Calca* ^104^ (Fig. S2A-B). Tc1 expressed *Cd4* (Fig. 8E) and showed high *S1pr1* and *Klf2* expression (Fig. 8C) consistent with a low activation and differentiation state ^105,106^. Tc2 expressed features of cytotoxic T cells including *CD8b1* and several killer cell lectin-like receptors (*Klrd1, Klra7, Klrg7, Klre1*) suggesting that the cluster was composed of conventional CD8 and natural killer T (NKT) cells. Cluster Tc3 was defined by a ISG signature (*Ifit3, Isg15, Rsad2, Usp18* among others) and included CD8 as well as CD4 T cells (Fig. 8C-E). This cluster was expanded at D14 after stroke indicating increased IFN signaling during the subacute phase of ischemic injury (Fig. 8B). Of note, CellChat analysis showed Tc2 and Tc3 cells interacting with Mg5, Mg6, BAM2, MdC4 and DC6 through *Cxcr3-Cxcl10* pathway at D14, indicating that myeloid *Cxcl10* expressing cells might have a role in the recruitment of CD8^+^ T cells into the brain ^107^ (Fig. 8F). Tc4 expressed T cell receptors of the γδT cell lineage (*Trdc, Trdv4, Tcrg-C1*). Expression of *Il17a*, a cytokine that has been implicated in aggrevating stroke pathology ^108-110^, was confined to the Tc4 cluster suggesting that γδT and not Th17 are the major IL-17 producing T cells after ischemic brain injury. Cluster Tc5 showed features of regulatory T cells (Treg) including the expression of the canonical transcription factors *Foxp3* and *Ikzf2* (also known as HELIOS). Interestingly, cells in this cluster expressed TNF receptors *Tnfrsf4, Tnfrsf9* and *Tnfrsf18,* which have been found in non-lymphoid tissue Treg but not in lymphoid-tissue associated Treg ^111^, possibly indicating that brain associated Treg don’t originate from lymphoid organs such as lymph nodes or spleen, but rather are recruited from non-lymphoid tissues or develop locally as previously suggested^11^. Similar to Tc3 and consistent with previous studies addressing Treg kinetics after ischemic brain injury ^11,112^, Tc5 was expanded at D14 after stroke (Fig. 8A-B). Cluster Tc7 was characterized by expression of cell proliferation markers including *Mki67*, *Top2a* and *Birc5* consistent with the presence of in situ T cell proliferation which was however not changed by stroke (Fig. 8B-C). No major changes were observed in the longitudinal composition of blood Tc (Fig. S10) which could be divided into five clusters constituting naïve CD8 (Tc1), CD4 (Tc2), memory CD8/NKT (Tc3) that showed a similar gene expression pattern as the brain Tc2 cluster, ISG T cells (Tc4), and a cluster with high expression of *Lgals1*, *Rora*, and several *S100* genes (Tc5).

**Figure 8.**
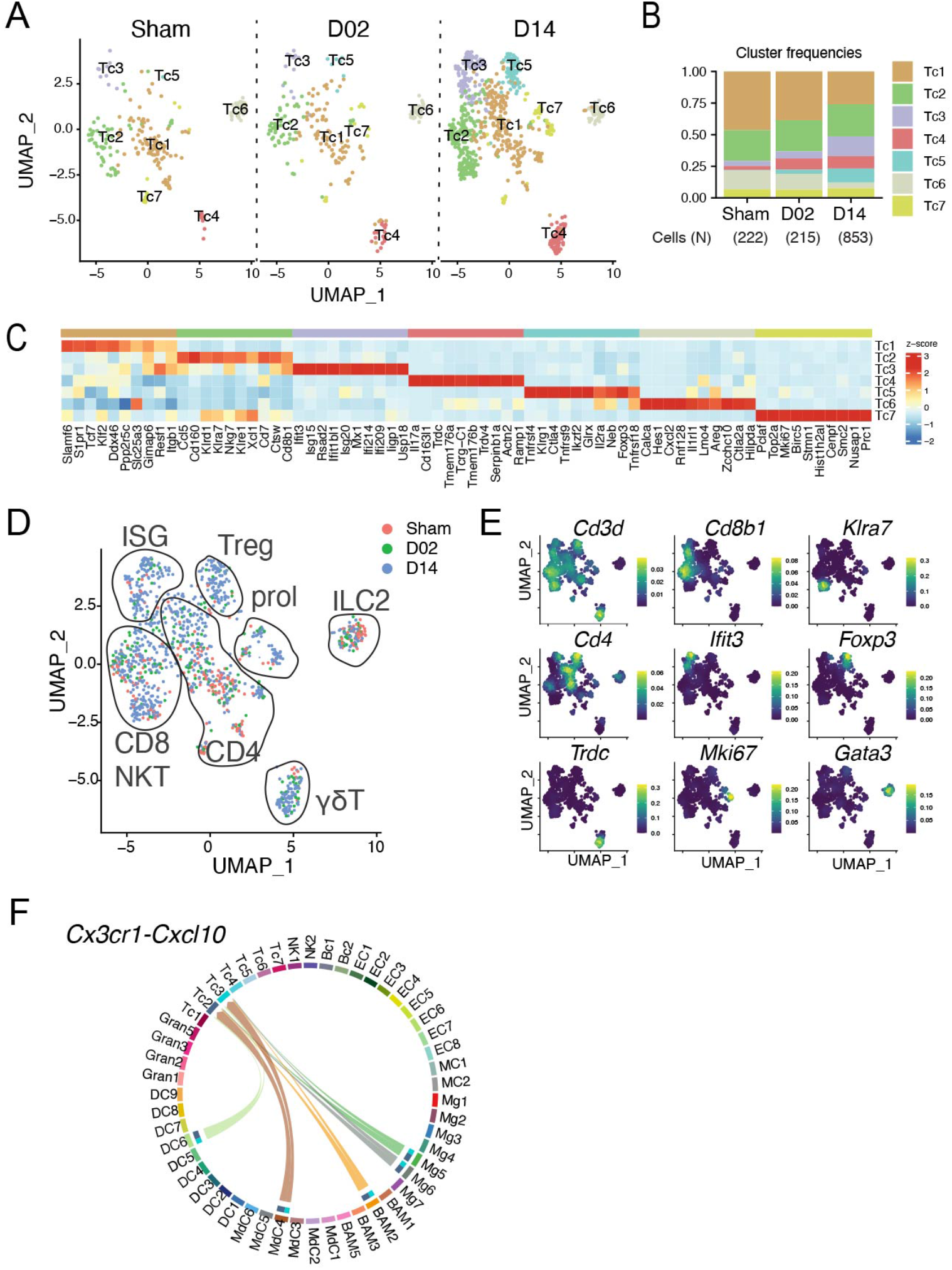
Transcriptional changes in brain lymphoid cells. **(A)** UMAP plots of brain lymphoid cells (Tc) transcriptomes for each studied time point identifies 7 clusters (Tc1-7). **(B)** Bar graph showing relative frequencies of Tc subclusters across Sham, D02 and D14 groups. **(C)** Heatmap displaying expression of the top 10 upregulated genes in each Tc subcluster. Scale bar represents Z-score of average gene expression (log). **(D)** UMAP of merged Sham, D02 and D14 Tc transcriptomes depicting identified Tc subtypes in diagrams: CD4, CD8, NKT, ILC2, ISG signature, γδT, Treg and proliferating T cells (prol). **(E)** Density plots in the UMAP space showing the expression of selected marker genes used for lymphoid cell type identification. Scale bar represents densities based on kernel density estimation of gene expression. **(F)** Chord plot showing cell-cell interactions between *Cxcr3* and *Cxcl10* in grouped Sham, D02 and D14 stroke mice. The strength of the interaction is indicated by the edge thickness. The color of the chord matches the cell cluster color sending the signal (*Cxcl10*). The number of cell cluster receptors (*Cxcr3*), and their weight in the interactions, is indicated by the color-matched stacked bar next to each cluster sender.

Similar to T cells, brain NK cells were increased at the D14 time point (Fig. S11B) and could be separated into two clusters (Fig. S11A). Brain NK1 was more similar to blood NK cells than brain NK2 as evidenced by higher Spearman correlation coefficient (Fig. S11D). CellChat analysis showed two major secretory axes confined to NK cells and some T cell, *Il18* and *Xcl1* signaling (Fig S11E-F). MdC4 were the main subset interacting with NK cells expressing *Il18r1* receptors, a known activator of NK cells ^113^, whereas clusters DC2 and DC9 interacted with NK cells via *Xcl1*-*Xcr1* pathway, possibly indicating a role of in cDC1 recruitment as previously reported ^114^.

Numbers and transcriptomes of brain associated B cells were not changed by stroke (Fig S11G) suggesting a minor role of B cells during the acute and subacute phases of ischemic brain injury.

### Brain transcriptomic changes after stroke in aged mice

The inflammatory response to ischemic brain injury differs between young and aged mice both in the transcriptomic response and the cellular composition of brain infiltrating immune cells ^115,116^. Therefore, we explored if aging alters cell signatures of immune and endothelial cells after stroke by preparing single cell transcriptomes from aged (17-20 months) Sham, D02, and D14 mice (Fig. S12A-E). The aged dataset was merged with the young brain dataset which was randomly down-sampled to match cell numbers of the aged dataset (aged = 11,999 cells). The combined data set showed distinct clusters for the major brain cell types including Mg, MdC, DC, Gran, Tc, BC, EC, and MC. The clusters of young and aged brains showed largely overlapping positioning in the UMAP space indicating the retention of the core transcriptome between the two groups (Fig. 9A). However, the frequency of infiltrating peripheral immune cell types differed between young and aged mice (Fig. 9C-D). Aged brains of Sham mice showed higher frequency of T cells while granulocytes and DC were more prominent in Sham brains of young mice indicating increased brain-associated T cells in brains of aged mice as previously reported ^117^. Overall, the cellular profile at D02 after stroke was comparable between groups, with a modest increase in neutrophils and decreased MdC in aged mice as previously reported ^118^ (Fig. 9C). At day 14, we observed increased Tc and reduced MdC and DC participation in aged as compared to young mice. At the subclass level we found that the distribution of Mg subclusters was similar between the age groups (Fig 9D). MdC clusters MdC5 and MdC6 were reduced in aged brains possibly reflecting the overall decreased brain MdC content at D14. Granulocytes showed a reduction in clusters Gran3 and Gran5 without overt changes in gene expression (Fig. 9D). T cells showed an expansion of CD8/NKT (Tc2) and ILC2 (Tc6) clusters whereas the frequency of γδT cells (Tc4) and Treg (Tc5) was reduced in aged brains ^119,120^. We analyzed differential gene expression of all cell clusters between aged and young mice and found that using a 2-fold change in expression level as a cutoff, overall, the number of genes that were exclusively either up- or downregulated in young stroke mice over young sham mice (D02 = 429 up, 380 down; D14 = 310 up, 406 down) was higher than in aged mice over aged sham mice (D02 = 73 up, 91 down; D14 = 67 up, 78 down) (Fig. 9E), whereas the majority of genes were regulated in both aged and young mice (Fig. 9E). We observed that young mice show higher number of regulated genes than aged mice, and that acute stroke (D02) led to higher exclusively regulated genes than subacute stroke (D14) in both young and aged mice (Fig. 9F). In addition, most of the exclusive DEG were detected in Mg, MdC, EC and to a lesser extend in Gran cell types (Fig. 9F). Among specifically upregulated genes in aged as compared to young mice, some belonged to the family of interferon inducible genes (*Ifi204*, *Cxcl10*, *Ifit3*, *Nfkbiz*) ^82^, which was reflected by a higher ISG score in several Mg, MdC, DC, and EC clusters in aged mice (Fig. 9G, Fig. S12F). Although it has been suggested that IFNβ signaling attenuates post-ischemic inflammation ^121^, the cellular sources of type I interferons in the post-ischemic brain have not been elucidated. In this study we found that *Ifnb1* was upregulated in some subsets of MdC and microglia in both young and aged mice (Fig. 9H).

**Figure 9.**
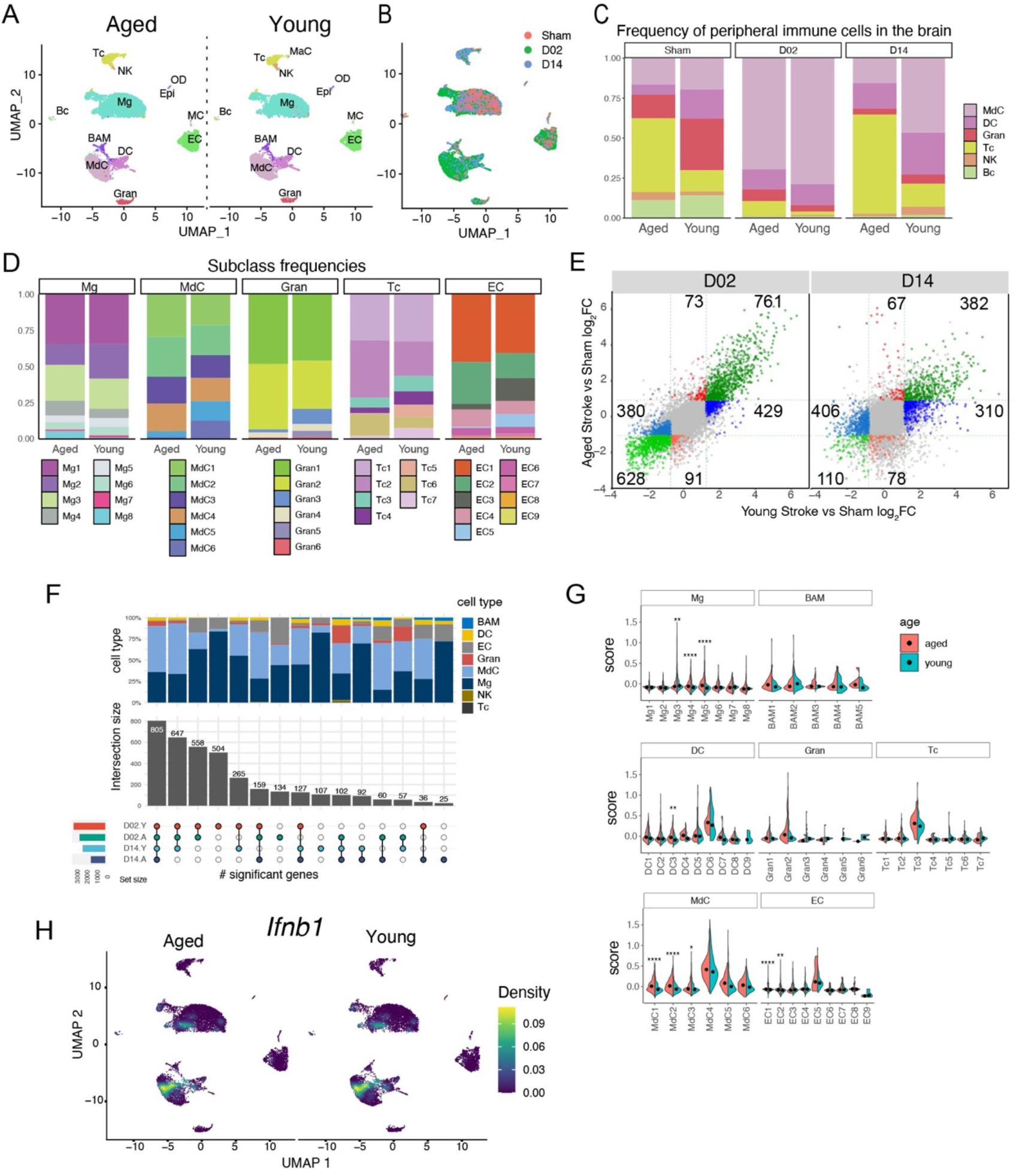
Comparison of the cellular composition and transcriptome signatures of brain and blood cells in aged and young stroke mice. **(A)** UMAP plot representing color-coded brain cell clusters identified in aged and young single-cell transcriptomes. **(B)** UMAP of 3 color-coded time point overlay of combined aged and young brain single-cell transcriptomes. **(C)** Bar graph of relative frequencies of infiltrating peripheral immune cells in the brain across Sham, D02 and D14 groups. **(D)** Bar graph showing relative frequencies of microglia (Mg), myeloid derived cells (MdC), granulocytes (Gran), T cells (Tc) and endothelial cells (EC) subclusters in either aged or young stroke mice. **(E)** Scatterplot comparing stroke-induced differential gene expression (DGE) versus Sham in young and aged mice at D02 and D14. Genes with log_2_FC > 1 are highlighted in color. Green color indicates differentially regulated genes in both aged and young mice. Blue color indicates differentially regulated genes in only young mice. Red color indicates differentially regulated genes in only aged mice. The number of differentially regulated genes in each subgroups is indicated in the corresponding subquadrants. **(F)** Upset plot of DGE results showing overlapping and age-specific (A, aged; Y, young) differentially expressed genes by cell type in response to either acute (D02) or subacute (D14) stroke. **(G)** Module-score analysis for interferon response gene sets in aged and young brain cell clusters. **(H)** Density plots in the UMAP space showing the expression of *Ifnb1* in the brain of young and age mice. Scale bar represents densities based on kernel density estimation of gene expression.

Together, the data provide a high-resolution landscape of the inflammatory response to ischemic brain injury in aged mice, which, compared to the response of young mice, is characterized by a different cellular composition but remarkable stability of the transcriptomic response at the single-cell level.

## Discussion

We sought to investigate the cellular immune landscape in the brain after transient cerebral ischemia during the early and late phases of the injury, and relate cellular signatures found in the brain to the cellular states of blood immune cells. In addition, we compared the cellular response between young and aged mice.

A major finding of this study is that there are distinct cellular transcriptomic responses of brain resident immune cells, infiltrating immune cells, and endothelial cells in the early and late phases of the tissue damage. Importantly, trajectory analysis showed that the transcriptome of blood-borne myeloid cells found in the brain after stroke remained distinct from their counterparts in the blood. These findings indicate that the local tissue milieu rather than peripheral immune priming determines the cellular state of monocyte-derived cells and neutrophils (Fig. 10).

**Figure 10.**
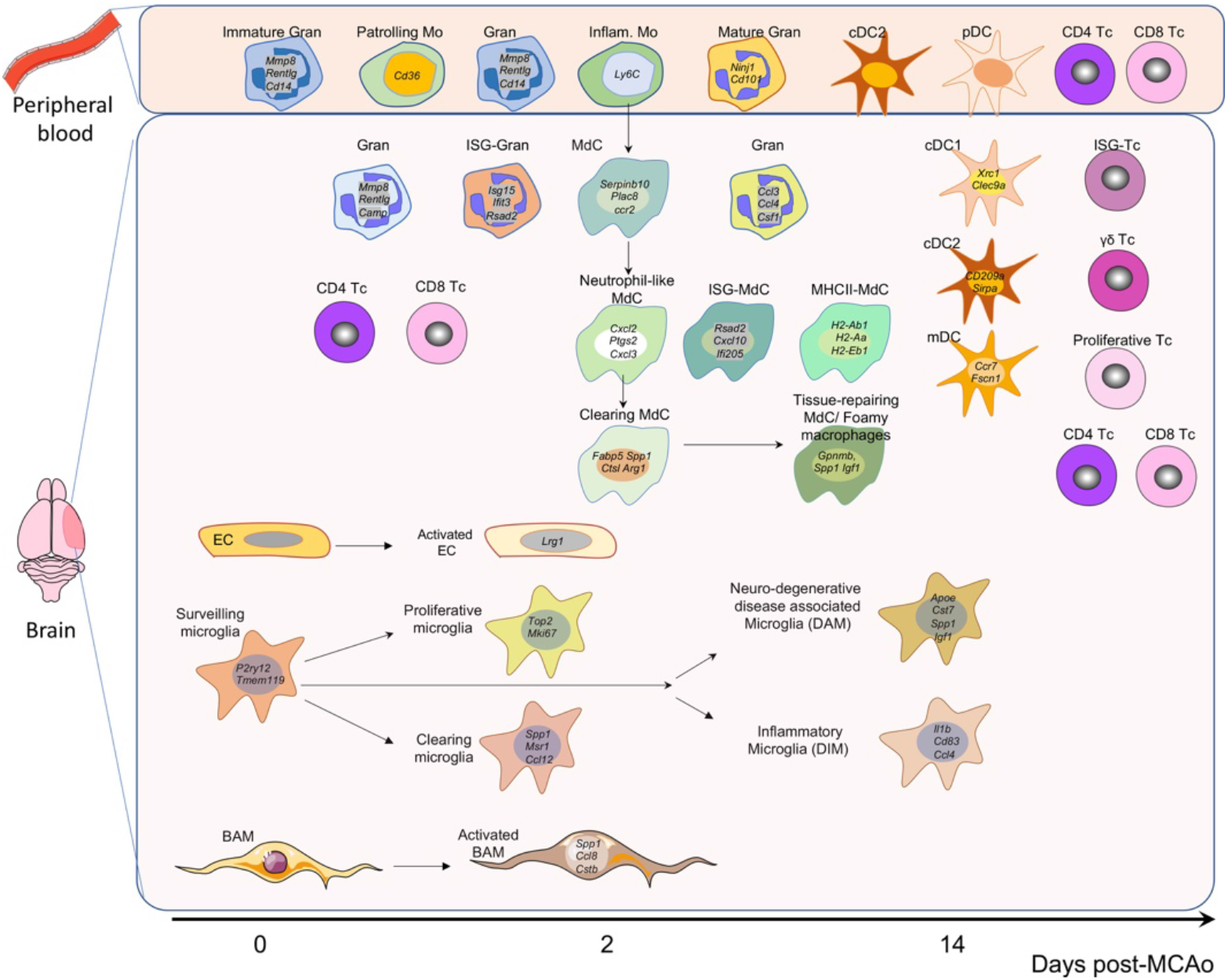
Summary of the main cell clusters identified in the blood and brain after experimental stroke.

Among the brain resident cells analyzed in this study, microglia showed the largest diversification in their transcriptional response across time points, whereas BAM and EC showed the strongest deviation from Sham states at D02 while the transcriptomes at D14 were more similar to Sham condition. Another key finding of this study is that microglia showed a strong proliferative response at D02 possibly triggered by the pronounced loss of microglia in the ischemic core. Similar proliferative responses have been observed after pharmacologic depletion of microglia ^122^, suggesting that depleting the microglial niche rather than a selective response to ischemic injury is inducing microglia proliferation. We found that proliferating microglia where primarily localized in the periinfarct region and that, at the D14 time point, the ischemic territory including the core areas was repopulated by microglia at a higher density than the one observed in non-injured tissue. Microglia acquired transcriptional states that were specific for the time points analyzed while there was no clear trajectory of microglia transcriptional states that would indicate a transition from the proliferative state observed at D02 to the inflammatory state that is prominent at D14.

Upregulation of interferon-stimulated genes has been observed in a variety of brain disease models including stroke ^47,48,123^. Similar to a previous study that identified ISG signatures in Mg, MdC, BAM, neutrophils, EC, and vascular smooth muscle cells one day after stroke ^17^, our study found ISG signatures in the same cell types but also in DC and Tc, suggesting a generalized IFN type I response independent of cell types and post-stroke time points (Fig. 10). Consistent with previous studies utilizing bulk RNA sequencing ^124^, we found a marginal but significant increase of ISG expression in aged animals after stroke, although the cell types and clusters showing this response were not changed. The IFN type I response is initiated by IFNα or IFNβ signaling, and we found *Ifnb1* expression primarily in MdC and to a lesser extend in Mg with a similar pattern in young and aged mice whereas we could not detect the expression of any of the *Ifna* gene subtypes (14 genes in the mouse).

MdC showed a continuous pseudotime trajectory originating from *Ccr2^hi^*/*Ly6c2^hi^*blood monocytes, gradually loosing monocytic marker genes and acquiring transcriptomic characteristics of tissue macrophages (Fig. 10). This is consistent with a longitudinal in situ transformation of macrophage phenotypes that is independent of de novo recruitment of blood monocytes as previously suggested ^125,126^. In contrast, and consistent with their short life span, brain granulocytes did not exhibit a pseudotime trajectory that was independent of their blood origins resulting in trajectories that suggest continuous recruitment from the circulating pool.

Another finding of this study is that cellular states of different myeloid populations can converge into similar transcriptomic phenotypes. For example, “monocyte derived-DC” (MdC5) showed high resemblance to DC clusters in their expression of MHC class II and co-stimulatory molecules while retaining an overall monocyte/MdC phenotype (expression of *Ly6c2, Lyz2, Cd14, Ms4a7*). Similarly, Mg clusters Mg3 and Mg7 share marker genes (*Spp1, Gpnmb, Apoe, Igf1*) with MdC6. In addition, we observed that microglia and MdC expressing the ISG marker gene *Cxcl10* were arranged into discrete clusters possibly indicating that myeloid cells organize into functional units to execute specific immune programs independently of their ontogeny.

A major limitation of this study is the restriction to two time points after cerebral ischemia. Our analysis at D02 and D14 identified high plasticity in the immune response and the transition of cell states can be only partially modeled on the provided data. For example, we have no information on the fate of proliferating microglia at D02 as we could not detect gene trajectories that would indicate progression to Mg clusters at D14. This is most likely due to the fact that transitional states might be present at time points not analyzed in this study. Similarly, endothelial cells showed a response at D02 that indicated initiation of migratory and proliferative response, whereas their transcriptomic profile at D14 was similar to Sham EC. Given that endothelial cells surrounding the infarcted brain area start to proliferate as early as 12–24 hours following occlusion and, that the angiogenic response and formation of new vessels continued more than 21 days post-stroke ^127,128^, it is likely that this response is not covered by our data.

Although our scRNA-seq study is one of the first to incorporate young and aged animals, we were not able to include aged male mice for the 14 day time point due to the high mortality rate observed in this group after transient focal ischemia, a drawback that has also been reported by others ^120^

Finally, because our investigation only included immune and endothelial cells, the response of other cell types including neurons, astrocytes, oligodendroglia, and pericytes were not assessed. This limitation is pertinent to the overall study design where we prioritized in depth resolution of cellular states of low abundance cell types such as BAM over the coverage of all brain cell populations. As more single cell RNA-seq datasets from experimental stroke studies become available it might be possible to integrate data from studies that used untargeted and targeted approaches to deliver a more comprehensive landscape of the immune response after stroke.

Taken together, by analyzing the immune response in the brain and peripheral blood at single cell resolution during acute and sub-acute stages after stroke, this study identifies cell type and time point specific immune programs and contributes to the ongoing efforts to compile a longitudinal cellular map of the immune response after stroke.

## Methods

### Mice

All procedures were approved by the institutional animal care and use committee of Weill Cornell Medicine and were conducted in accordance with the ARRIVE guidelines ^129,130^. Experiments were performed in young (8-12 week-old) male wild type mice obtained from Jackson Laboratory (IMSR_JAX:000664; Bar Harbor, ME) and aged (17-18 month-old) male and female wild type mice obtained from the NIA-NIH. All in house bred mice were on a C57Bl/6J background and included B6.Cg-Gt(ROSA)26Sortm14(CAG-tdTomato)Hze/J (IMSR_JAX:007914), B6.129P2(C)-Cx3cr1tm2.1(cre/ERT2)Jung/J (IMSR_JAX:020940) and C57BL/6-Tg(CAG-EGFP)131Osb/LeySopJ (IMSR_JAX:006567).

### Middle Cerebral Artery occlusion

Transient focal cerebral ischemia was induced using the intraluminal filament model of middle cerebral artery (MCA) occlusion as described previously ^131^. Briefly, under isoflurane anesthesia (maintenance 1.5-2%), the MCA was occluded for 35 minutes using a 6-0 Doccol monofilament (#L12; Sharon, MA). Reperfusion was confirmed by measuring the cerebral blood flow (CBF) in the MCA territory by transcranial laser Doppler flowmetry (Periflux System 5010, Perimed, King Park, NY). Only animals with CBF reduction of >85% during MCA occlusion and CBF recovered by >80% after 10 minutes of reperfusion were included in the study (Jackman et al., 2011). Rectal temperature was monitored and kept constant (37.0±0.5°C) during the surgical procedure and in the recovery period until the animals regained full consciousness. After, mice were housed in a single chamber environmental incubator (Darwin Chambers, Saint Louis, MO) to hold mice at 29-32°C for up to 4 days. Topical lidocaine and bupivacaine (0.25%, 0.1 ml, transdermal) were used for pre-operative analgesia and buprenorphine (0.5 mg/kg q12h; s.c.) as post-operative analgesia for 72 hours.

### Tamoxifen treatment

Four-week old male Cx3cr1^CreER^-Td-tomato mice were treated with tamoxifen (T5648, Sigma; 80mg/Kg) dissolved in corn oil (C8267, Sigma) by intraperitoneal injection during 5 consecutive days. The mice were used for experiments 6 to 8 week after treatment.

### Cell isolation

Isolation of brain and blood cells from sham and stroke mice were performed at the same time of the day (8:00–9:00 am) for all experiments to limit the effects of circadian gene expression variation ^132^. Mice were anesthetized with pentobarbital (100 mg/Kg, i.p.) and transcardially perfused with heparinized PBS (2U/ml). Briefly, either control-sham or ischemic hemispheres were separated from the cerebellum and olfactory bulb and gently triturated using a Gentle MACS dissociator (Miltenyi Biotec, Auburn, CA). Single cell suspensions were obtained by enzymatic digestion with papain (Neural Tissue Dissociation Kit (P), #130-092-628, Miltenyi Biotec) following the manufacturer’s instructions. Four to five hemispheres were pooled for each experiment. In order to increase cell viability and to preserve the transcriptional state during the generation of single-cell suspensions, Brilliant Blue G (BBG, P2X7 receptor antagonist, 1μM, Sigma), AP-5 (25 nM, NMDA receptor blocker, Tocris), and actinomycin D (RNA polymerase inhibitor, 5μg/ml, Sigma) were added to the dissociation solution ^133^ ^134^. Next, cell suspension was filtered through a 70 μm filter, resuspended in 30% Percoll (GE Healthcare)-HBSS containing 1μM BBG, and spun for 10 minutes at 700 *g*. After gradient centrifugation, the myelin layer was removed, and the cell pellet was resuspended in 2%FBS-PBS buffer and filtered through a 40 μm filter. Isolated cells were washed and resuspended in 100 μl of blocking buffer per hemisphere to proceed for FACS staining and cell sorting.

For isolation of peripheral leukocytes, mice were anesthetized with pentobarbital (100 mg/Kg, i.p.) and 0.5 ml of blood was collected by cardiac puncture into heparinized tubes. For each experiment the blood from two mice (1 ml total blood) was pooled and erythrocytes were lysed. BBG (1μM) and actinomycin D (5μg/ml) inhibitors were added during the isolation procedure. After erythrolysis, blood cells were resuspended in MACS buffer (PBS supplemented with 2% FBS, 2 mM EDTA; 300 μl/10^7^ cells) and incubated with a biotinylated Ter-119 antibody (Table S1) and remaining erythrocytes were depleted with anti-biotin microbeads according to the manufacturer’s instructions (Miltenyi Biotec). Afterward, cells were washed and resuspended in 0.01% BSA-PBS at a concentration of 10^5^ leukocytes/ml for Drop-seq processing.

### Flow Cytometry and Cell Sorting

For Drop-seq experiments, brain single cell suspensions were incubated with anti-CD16/CD32 antibody for 10 min at 4°C to block Fc receptors, followed by staining with CD45-BV510, Ly6C-FITC, CX3CR1-PE, Ly6G-PercP-Cy5.5 and CD11b-APC-Cy7 antibodies for 15 min at 4°C (Table S1). CD45^hi^ cells, microglia (CD45^hi^CD11b^+^CX3CR1^+^) and endothelial cells (CD45^-^Ly6C^+^) were sorted on an Aria II cytometer (BD Bioscience) and collected in 0.5 ml of 0.01% BSA-PBS for Drop-seq. Flow cytometry gating strategy is described in Fig. S1. Analytical flow cytometry was performed on a NovoCyte Flow Cytometer (Agilent, Santa Clara, CA). The antibodies used are described in Table S1. Appropriate isotype controls, ‘fluorescence minus one’ staining, and staining of negative populations were used to establish gating parameters.

### Generation of single cell RNA libraries by Drop-seq

Single-cell transcriptomes of sorted brain cells and purified blood leukocytes were prepared by drop-seq as described ^135^ with modifications. Cells were resuspended in PBS-0.01%BSA to a final concentration of 100 cells/μl. Barcoded capture beads (ChemGenes Corporation, Wilmington, MA) were resuspended in 1.8 ml lysis buffer consisting of 4M Guanidine HCL (Thermofisher Scientific, Waltham, MA), 6% Ficoll PM-400 (Sigma-Aldrich), 0.2% Sarkosyl (Sigma-Aldrich, St. Louis, MO), 20mM EDTA (Thermofisher Scientific), 200mM Tris pH 7.5 (Sigma-Aldrich), 50mM DTT (Sigma-Aldrich) at a concentration of 120 beads/μl. A 5mm diameter, 1.7mm thick PVDF encapsulated magnetic stir disc and rotary magnetic tumble stirrer (V&P Scientific, San Diego, CA) was used along with a 3 ml syringe that contained the beads in lysis buffer to keep the beads in suspension. Thereafter, single cells and beads were encapsulated in nanoliter-scale droplets using a Drop-seq microfluidic device coated with Aquapel (FlowJEM, Toronto Canada), droplet generation oil (BioRad, Hercules, CA) using flow rates of 4 ml/h for cells and beads and 15 ml/h for oil. Each run typically lasted about 18 minutes. After removing the oil, droplets were resuspended in 30 ml of room temperature 6X SSC (Promega, Madison, WI) and 1 ml perfluorooctanol (Sigma-Aldrich, St. Louis, MO) and shaken vigorously 6 times vertically to break the droplets. The beads were captured by loading them into a 20 ml syringe with an attached 0.22µm Millex-Gv syringe filter (Millipore Sigma, Burlington, MA) as previously described ^136^. Beads were washed with 2 x 20 ml of ice cold 6X SCC. The syringe filter was then inverted, and a 10 ml syringe was used to flush the beads out with 10 ml ice cold 6X SSC repeated for a total of 3 times. Beads were collected by centrifugation at 1,250 x g for 2 min at 4°C with low brake setting. The remainder of the Drop-seq protocol followed the one published ^135^. cDNA was amplified by PCR using the following parameters: 95°C (3 min); 4 cycles of 98°C (20 s), 65°C (45 s), 72°C (3 min); 11 cycles of 98°C (20 s), 67°C (20 s), 72°C (3 min). Libraries were quantified by quantitative PCR and checked for quality and size distribution on a Bioanalyzer (Agilent) before sequencing on an Illumina NextSeq500 instrument using the 75 cycle High Output v2 kit (Genomics Core Facility, Cornell University, Ithaca, NY). Three to four libraries were multiplexed into a single run. We loaded 1.8 pM library and provided Drop-seq Custom Read1 Primer at 0.3 μM in position 7 of the reagent cartridge without PhiX spike-in using a read configuration of 20 bases (Read1), 8 bases (Index1), and 64 bases (Read2). Details regarding each Drop-seq run and characteristics of the strain, sex, age, and number of mice used for DropSeq can be found in Table S2.

### Data pre-processing

Demultiplexed fastq files were cleaned of reads not passing the Illumina Passing Filter with fastq_illumina_filter (version 0.1) and processed with the Drop-seq Tools (version 2.3.0) pipeline ^137^. Briefly, each transcriptome Read2 was tagged with the cell barcode (bases 1 to 12) and unique molecular identifier (UMI) barcode (bases 13 to 20) obtained from Read1, trimmed for sequencing adapters and poly-A sequences, and aligned using STAR v2.7.3a ^138^ to the mouse reference genome assembly (Ensembl GRCm38.94 release). Reads aligning to exons were tagged with the respective gene symbol and counts of UMI-deduplicated reads per gene within each singular cell barcode were used to build a digital gene expression (DGE) matrix. The DGE matrix contained 40,000 cell barcodes associated with the highest numbers of UMIs. We used the DecontX method from the R package celda ^139^, a Bayesian hierarchical model to estimate and remove cross-contamination from ambient RNA, to construct a corrected DGE matrix. Cells with fewer than 200 UMIs, more than 10,000 UMIs, or more than 20% mitochondrial genes were excluded. We used Doubletfinder (RRID:SCR_018771) ^140^ to computationally detect cell doublets with an expected doublet rate of 5% ^135^ as input parameter. Cells tagged with a “Doublet” call were removed. Finally, the corrected DGE matrices were merged into a single matrix.

### Bioinformatic analysis and statistics

We used Seurat (version 4.1.0; RRID:SCR_016341) ^141^ for down-stream analysis.

1. Counts were log-normalized for each cell using the natural logarithm of 1 + counts per ten thousand.
2. The 3000 most variable genes were identified by calling *FindVariableFeatures*.
3. We next standardized expression values for each gene across all cells by Z-score transformation (*ScaleData*).
4. Principle Component Analysis (PCA) was performed on the scaled variable gene matrix. The R package Harmony (RRID:SCR_022206) ^142^ was used to correct the matrix for batch effects. The data from replicate experiments were combined into four sets so that each set included a Sham, D02, and D14 experiment. We run Harmony on the first 40 PCA dimensions with a maximum of 20 iterations.
5. We used the Uniform Manifold Approximation and Projection (UMAP) ^143^ for dimensional reduction and visualization of Harmony-derived embeddings in a two-dimensional space with preset parameters by invoking the *RunUMAP* function in Seurat utilizing the 40 first components of the Harmony reduction.
6. We used the Louvain algorithm as implemented in *FindClusters* with a resolution setting of 1.2 for the brain and 0.7 for the peripheral blood dataset to perform graph-based clustering on the neighbor graph that was constructed with the *FindNeighbors* function call on Harmony-derived embeddings.
7. After clustering, we used the model-based analysis of single cell transcriptomics (MAST) algorithm ^144^ in the *FindAllMarkers* function to find differentially expressed genes in each cluster based on the log-normalized expression matrix with parameters: only.pos = T, min.pct = 0.1, logfc.threshold = log2(1.5), max.cells.per.ident=2000.

We performed unsupervised cell type annotation using the SingleR package (RRID:SCR_023120) ^145^ with ImmGen ^146^, BrainImmuneAtlas ^55^, and Tabula Muris ^147^ as reference datasets (see Fig. S2). Assignments were further manually validated by scoring the 10 most differentially expressed genes for the presence of canonical marker genes for each cell type. On these bases, we assigned the metacells to microglia (Mg), border-associated macrophages (BAM), monocyte-derived cells (MdC), granulocytes (Gran), mast cells (MaC), dendritic cells (DC), T cells (Tc), NK cells (NK), B cells (Bc), endothelial cells (EC), vascular mural cells (MC), epithelial-like cells (Epi), and oligodendrocytes (OD) clusters for the brain dataset, and monocytes (Mo), granulocytes (Gran), eosinophils/basophils (Eos/Bas), dendritic cells (DC), T cells (Tc), NK cells (NK), B cells (Bc), various precursors (pre), and one unclassified cluster (UC) for the peripheral blood dataset.

To achieve further resolution of cell states, individual count matrices were generated based on the initial cluster designation and steps 1-7 were repeated with the following modifications: Mitochondrial, ribosomal, and gene model (*Gm*) annotated genes were removed; Step 2, 2000 variable features were selected; Step 4, Harmony was run on the first 15 PCA dimensions; Step 5, The “min.dist” parameter in the *RunUMAP* function was set to 0.1 for clusters with more than 2500 cells; Step 6, *FindClusters* was performed at a resolution of 0.4. Differentially expressed genes with an FDR < 0.05 were ranked by their log2 fold change and z-scores were computed on the average gene expression across clusters for visualization in heatmaps.

### Cluster pruning and metacell exclusion

After subclustering we detected several clusters within the brain dataset with high expression of microglial (*Hexb, Siglech, P2ry12*) marker genes together with either granulocyte (*S100a8, S100a9, Cxcr2*), DC (*Cd209a, Xcr1, Ccr7*), T cells (*Trac, Trbc2, Cd3d*), NK cells (*Nkg7, Gzma*), and macrophage (*Mrc1, Lyve1*) genes. Although we cannot exclude that this is due to biological processes such as transcriptomic changes or engulfment of living cells by microglia as previously reported for neutrophils and lymphocytes ^148^, we opted to exclude these cells as potential cell doublets. Therefore, we manually removed 1730 Mg, 287 DC, 145 BAM, 70 Gran, 65 Tc, and 41 NK metacells. After exclusion of these cells, we rerun steps 1-7 on the main dataset and performed de novo subclustering as described above.

### Analysis of combined young and aged brain datasets

The dataset from aged (17-20 month) mice brains were preprocessed as described above. Cell identities in the aged mice brain dataset were assigned using the *FindTransferAnchors* and *TransferData* functions in Seurat using the young brain dataset as reference. Seurat objects from young and aged mice were merged into a single object by retaining all cells of the aged brains and randomly down-sampling the young brain dataset. This resulted in balanced cell numbers among cell type classes in both datasets (Table S3). Raw count data were processed as described above. The Harmony algorithm was run on the first 40 principal components of the PCA. UMAP was computed on the first 40 Harmony dimensions. Differential gene expression (Fig 9E) was computed with limma-voom with default parameters ^149^ after “pseudobulk” conversion of individual experiments using the *aggregateAcrossCells* function of the scuttle package ^150^ following the removal of mitochondrial, ribosomal, hemoglobin, gene model (Gm) annotated genes, and the sex-specific genes (*Tsix, Xist*).

### Cell trajectory inference

We conducted the trajectory inference (TI) analysis using the Dyno package ^151^. Brain and peripheral blood datasets were merged and randomly downsampled to a maximum of 1000 cells per cluster and treatment. The merged dataset was subset to contain either Mg, blood Mo and brain MdC, or blood Neu and brain Gran clusters. The TI was performed on the 2000 most variable genes selected with Seurat *FindVariableFeatures* function. For Gran and Mo/MdC, blood granulocytes and monocytes were designated as starting population, respectively. For Mg, the starting population consisted of all Mg from Sham brains. The most appropriate TI method (Slingshot) ^152^ was selected based on Dyno (*guidelines_shiny* function) recommendations. UMAP was calculated on the first 15 PCA dimensions and the 2-dimensional space and trajectories were visualized by the *plot_dimred* function of the Dyno package.

### CellChat analysis

Cell-cell interaction networks were constructed using the CellChat R package (RRID:SCR_021946) ^153^ with a custom mouse ligand-receptor interaction database that contained combined entries of the curated RNAMagnet ^154^ and CellChat databases. Briefly, the processed Seurat object was split by treatment into Sham, D02, and D14 datasets. Then, cell clusters were randomly downsampled to contain no more than 1000 cells. The matrices were used as input for the *createCellChat* function and processed using its standard pipeline. Differentially expressed genes and interactions were identified in the CellChat object via *identifyOverExpressedGenes* and *identifyOverExpressedInteractions*, respectively. The CellChat algorithm was then run to calculate the probable interactions at the cell-to-cell level via *computeCommunProb* with “truncatedMean” as the method for computing the average gene expression after removing 15% of observations from each end of the gene expression vector (parameter: trim = 0.15). The *filterCommunication* function was used to filter out interactions with less than 20 cells in each cluster. Communication probabilities on the signaling pathway level were then calculated by invoking the *computeCommunProbPathway* function. Ligand-receptor interaction probabilities were visualized via *netVisual_aggregat*e and *netVisual_individual*. For joint analysis of different group datasets (Sham, D02, D14), CellChat objects were merged and *compareInteractions* was used to compare the total number and strength of interactions between treatments followed by the *rankNet* function for visualization.

### Modular score calculation and visualization

*AddModuleScore* function in the Seurat package was used to calculate the functional signatures of each cell cluster. The type I interferon-response score was calculated using interferon-stimulated genes (ISG) as previously reported ^155^. Disease-associated microglia (DAM) and disease inflammatory macrophages (DIM) scores were calculated using the gene list reported by Silvin et al.^38^. Additional module scores were calculated for Foamy Macrophages ^63^, Stroke-Associated macrophages (SAM) ^16^, and monocytes ^156^. All marker genes used for score calculation can be found in Supplementary Table S4. Module scores were visualized in R using the ggplot2 (V3.3.6) package (RRID:SCR_014601) ^157^.

### Correlation plots

Cluster-wise average gene expression was calculated with the *AverageExpression* function of the Seurat package. The *cor* function in R were used to construct a Spearman correlation matrix of gene expression between blood monocyte and brain MdC clusters. The correlation matrix was visualized with the *corrplot* function of the corrplot (V0.92) package (RRID:SCR_023081) ^158^.

### Combined analysis of young and aged brain scRNA-seq data

Cell identities were assigned by aligning both datasets in the high-dimensional space and projecting cluster annotations from the young dataset onto the aged dataset essentially as previously described ^141^ and by using SingleR to adjudicate cell identities using the young dataset as a reference. For each cell the maximal correlation score obtained by the two methods was used to assign the cell identity. The young dataset was down-sampled to contain no more than 5000 cells per treatment and both datasets were merged and processed using steps 1-6 as outlined above (Bioinformatics analysis). Differential gene expression between aged and young mice was calculated for each cell cluster using the MAST algorithm of the *FindMarkers* function in Seurat with preset parameters after excluding mitochondrial, ribosomal, hemoglobin, sex-specific (*Tsix*, *Xist*), and gene model (Gm) annotated genes. Genes were considered differentially regulated if they showed higher than two-fold change in expression level and an FDR < 0.05.

### Brain histology

Mice were deeply anesthetized with sodium pentobarbital and transcardially perfused with ice cold PBS (30ml) followed by 4% paraformaldehyde (PFA) in PBS (100ml). Brains were dissected, post-fixed in 4% PFA overnight, dehydrated in 30% PBS-sucrose solution for 1-2 days and then frozen using dry ice. Frozen brains were then embedded in Epredia™ M-1 Embedding Matrix (Thermo Ficher Scientific) and cut in coronal sections using a cryostat (Leica CM3050S, Mannheim, Germany), and mounted on slides for either fluorescence immunocytochemistry (IF) downstream applications or RNA-fluorescence in situ hybridization (FISH).

For IF, coronal 18 μm-thick sections were permeabilized with 0.5% Triton X-100 (Sigma) in PBS (PBST), blocked with 5% normal donkey serum (NDS) in 0.1% PSBT for 1 hour and incubated overnight at 4°C with primary antibodies (table 2) in 1%NDS-0.1% PBST. After ON incubation, sections were washed 3×5min with 0.1%PBST and incubated with secondary antibodies in 1%NDS-0.1%PBST for 1 hour at RT. Sections were washed with 0.1% PBST 2×5min, followed by a 1×5min wash in 0.1%PBST-DAPI (1:100, 12.5ng/ul), mounted with FluorSave Reagent (Millipore) and visualized using either the Olympus IX83 or Leica TCS SP8 confocal microscopes. Primary and secondary antibodies used are described in Table S5.

RNA-FISH was performed using RNAscope Multiplex Fluorescent Kit v2 (ACD-Bio-Techne, Newark, CA) following the manufacturer’s instructions. Briefly, 10-μm thickness sections were processed using RNAscope hydrogen peroxide, followed by boiling Target Retrieval solution, dehydrated by 100% ethanol, and incubated in Protease III solution. Afterwards, tissue sections were hybridized with the target probes (Table S6) for 2 h at 40 °C, followed by a series of signal amplification and washing steps. Hybridization signals were detected by fluorescent signal using peroxidase-based Tyramide Signal Amplification-Plus Fluorescein or TSA Plus Cyanine 5 (Perkinelmer, Shelton, CT). Finally, the sections were counterstained with DAPI and coverslipped using ProLong Gold Antifade Mountant. Images were acquired using either an epi-fluorescent (Olympus IX83, Waltham, MA) or a confocal (Leica) microscope. Images were analyzed using Fiji (RRID:SCR_002285) ^159^. When FISH and IF staining were combined, the slides were washed in RNAscope wash buffer after the development of Tyramide Signal and then, IF was sequentially performed as described above but skipping the permeabilization step.

## Data availability

The raw and processed data and metadata of all scRNA-seq datasets included in this study are available in the GEO repository (GSE225948). A public accessible interactive web portal for exploring the scRNA-seq data included in this study has been developed (https://anratherlab.shinyapps.io/strokevis/).

## Acknowledgments

We thank Dr. Christopher Mason for helpful discussions. The generous support of the Feil Family Foundation is gratefully acknowledged.

## Author contributions

J.A and L.G-B conceived the study with input from C.I; L.G-B, Z.S, R.S, O.N and G.R performed experiments and analyzed data; JA performed bioinformatic analyses. L.G-B and J.A wrote the original draft. C.I revised the manuscript. All authors read and approved the final manuscript.

