## Supplementary figures for "Brain and blood single-cell transcriptomics in acute and subacute phases after experimental stroke"

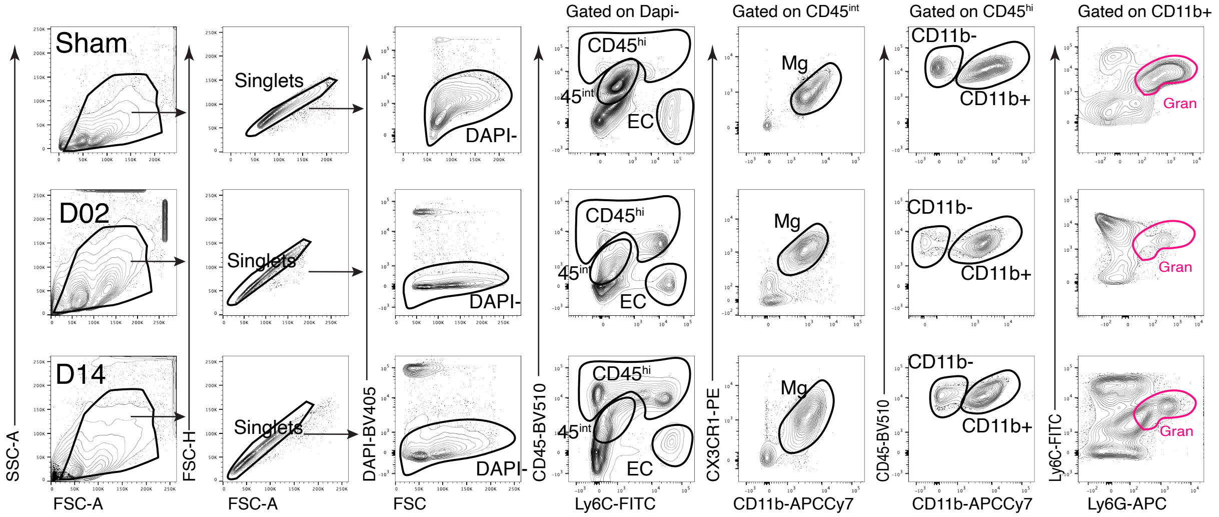


**Figure S1. Flow Cytometry strategy for sorting brain single cell suspensions for DropSeq.** Representative flow cytometry plots of the gating scheme used to identify and sort CD45hi cells, endothelial cells (EC) and microglia from Sham, D02 and D14 single cell suspensions of mouse brains. Live cells (DAPI–) were gated based on forward and side scatter and DAPI staining. CD45hi and ECs (CD45loLy6Chi) were sorting based on CD45 and Ly6C expression. CD45int was selected for further analysis and examined for the expression of CD11b and CX3CR1 to identify microglia (CD45intCD11b+CX3CR1+). Analysis of CD11b and Ly6G expression was assessed in CD45hi subpopulation to verify the presence of granulocytes (CD11b+\ Ly6G+), other myeloid cells (CD11b+\ Ly6G–) and lymphocytes (CD11b–).

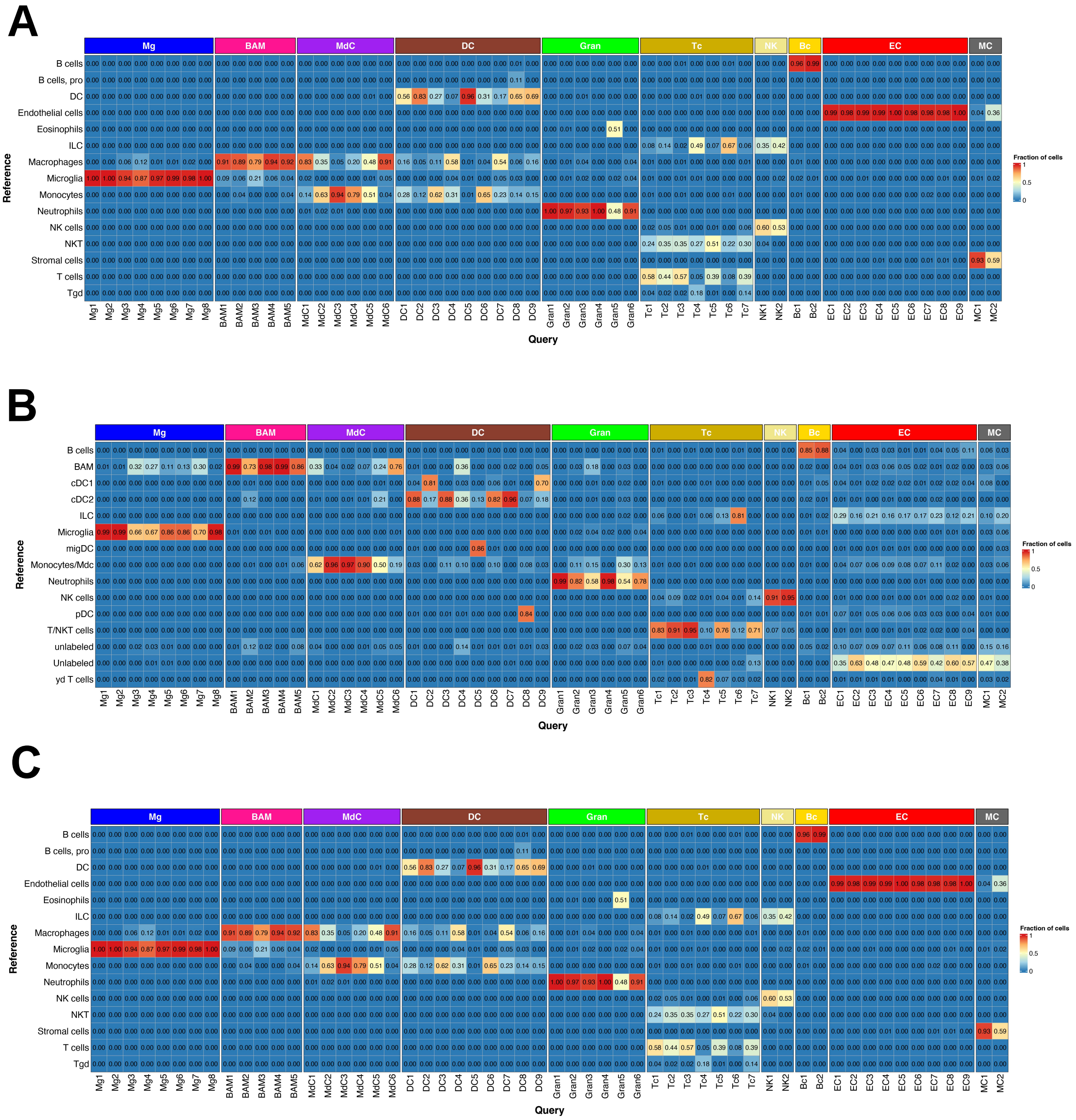


**Figure S2. Unsupervised cell annotation of the brain dataset.** Heatmaps showing the percentage of cell identity of each cluster (columns) attributed to a cell type classifier (rows) after unsupervised annotation with Immgen (**A**), BrainImmuneAtlas (**B**) or Tabula Muris (**C**) reference datasets using SingleR.

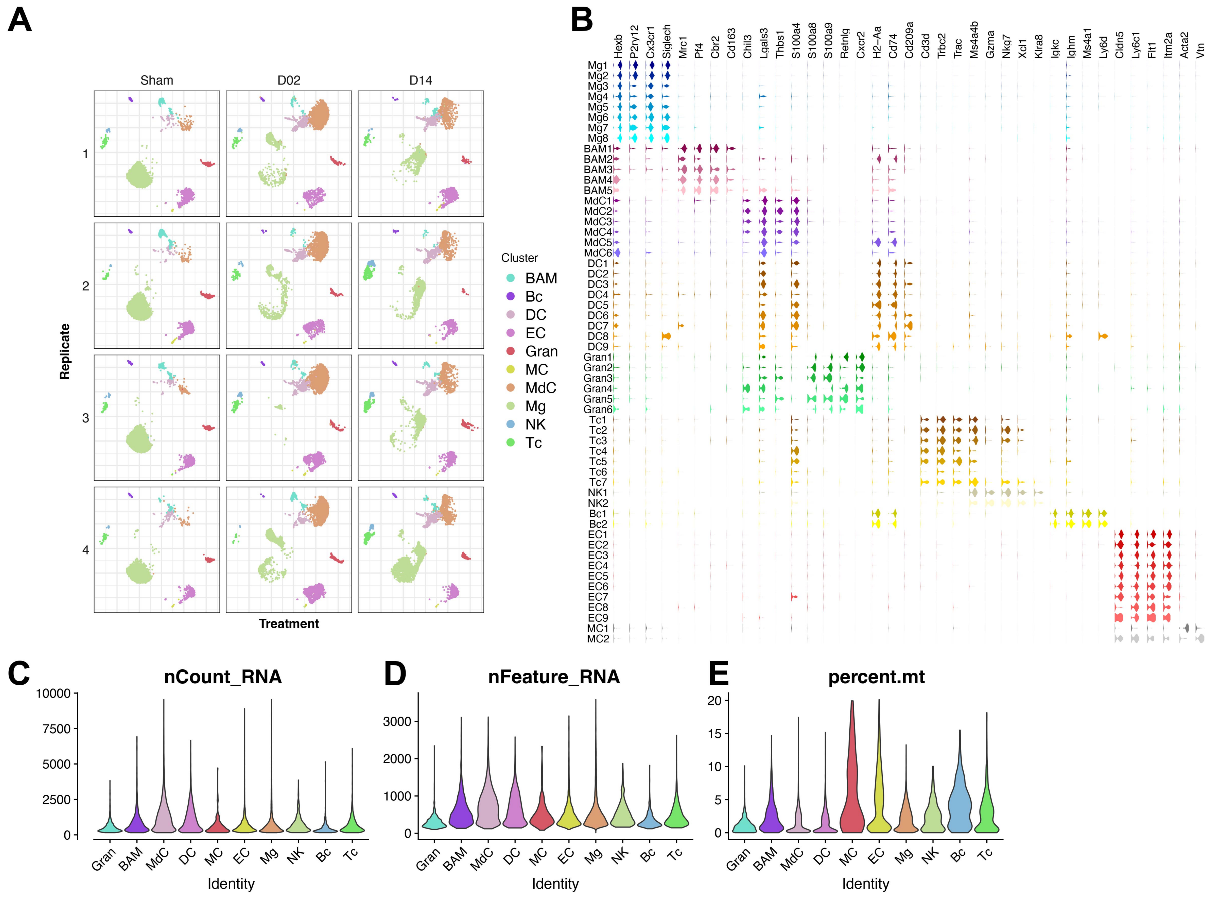


**Figure S3. Brain cell transcriptomic data set quality**

**(A)** UMAPs of the identified clusters for each biological replicate of brain cell transcriptomes from control surgery (Sham) or stroke mice 2 (D02) or 14 (D14) days after injury.

**(B)** Violin plots generated from the integrated dataset displaying characteristic cell marker genes of each identified cell subcluster.

**(C)** Violin plots showing distribution of number of total UMI counts per cell (nCount), genes detected per cell (nFeature), and percentage of mitochondrial genes (percent.mt) per identified cell type. Microglia (Mg), border-associated macrophages (BAM), monocyte-derived cells (MdC), granulocytes (Gran), mast cells (MaC), dendritic cells (DC), T cells (Tc), NK cells (NK), B cells (Bc), endothelial cells (EC), vascular mural cells (MC).

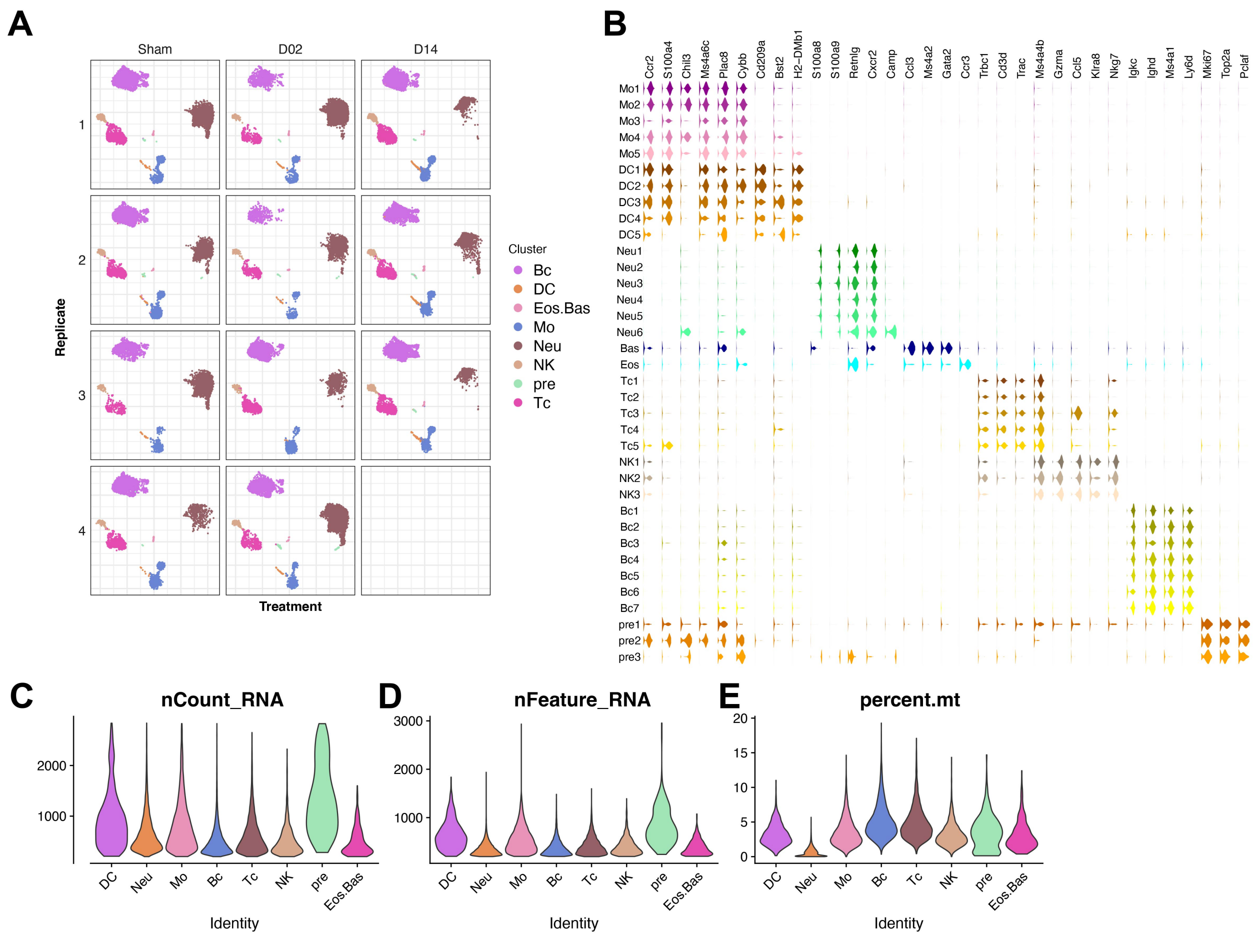

**Figure S4. Blood cell transcriptomic data set quality**

**(A)** UMAPs of the identified clusters for each biological replicates of peripheral blood cell transcriptomes from control surgery (Sham) or stroke mice 2 (D02) or 14 (D14) days after injury.
**(B)** Violin plots generated from the integrated dataset displaying characteristic cell marker genes of each identified cell subcluster.
**(C)** Violin plots showing distribution of number of total UMI counts per cell (nCount), genes detected per cell (nFeature), and percentage of mitochondrial genes (percent.mt) per identified cell type. Monocytes (Mo), granulocytes (Gran), eosinophils/basophils (Eos/Bas), dendritic cells (DC), T cells (Tc), NK cells (NK), B cells (Bc), hematopoietic precursors (pre).

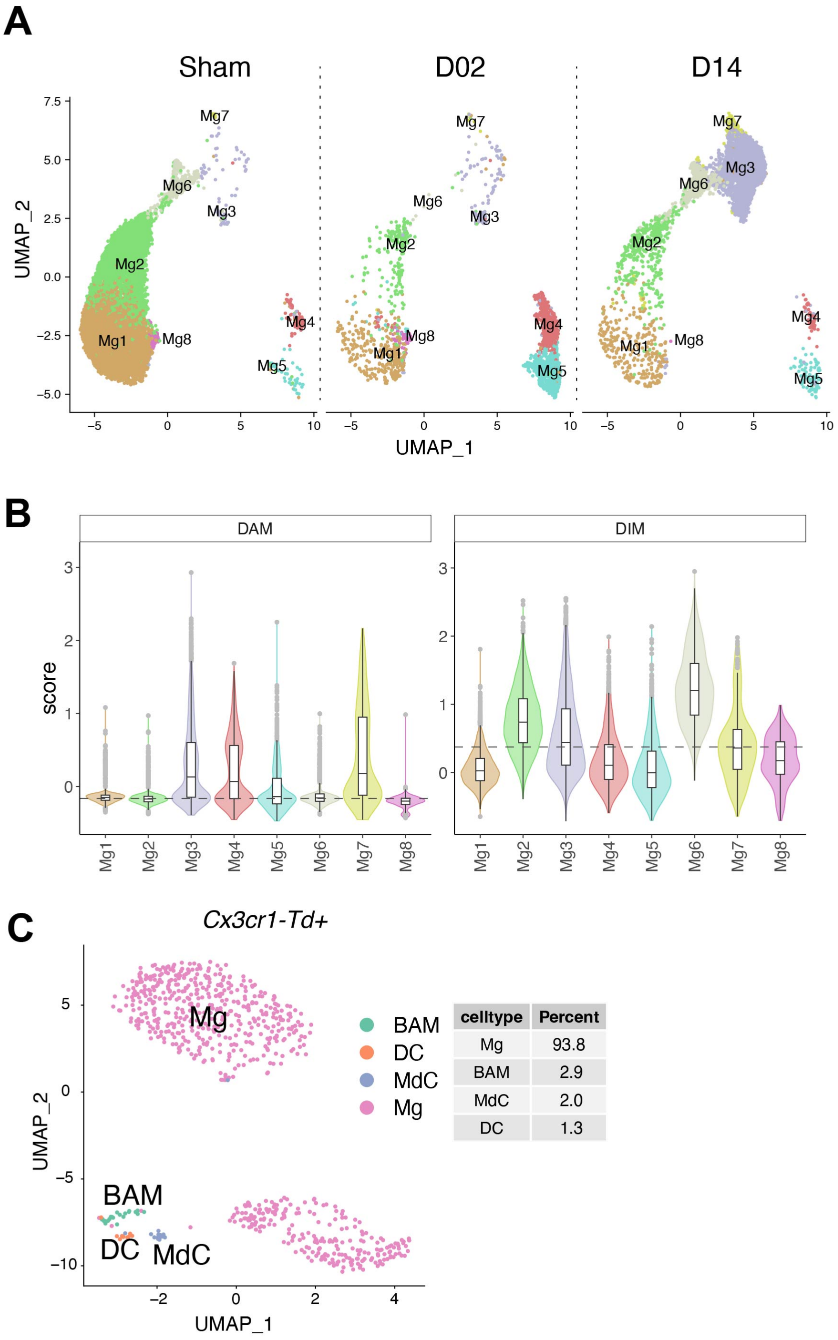


| celltype | Percent |
|---|---|
| Mg | 93.8 |
| BAM | 2.9 |
| MdC | 2.0 |
| DC | 1.3 |

**Figure S5. Microglia clusters across conditions and module score analysis**

**(A)** UMAP plot showing identified subclusters of microglial cells (Mg) split by studied time points: control surgery (Sham) or stroke mice 2 (D02) or 14 (D14) days after injury.
**(B)** Violin plots depicting module score for disease-associated microglia (DAM) and disease inflammatory macrophages (DIM) of brain microglia clusters. Dashed line shows the average score across all Mg.
**(C)** UMAP plot of *Cx3cr1* -Td+ brain cell transcriptomes (Table 2) showing most of the Td+ cells annotated as microglia (93.8%). The analysis was performed using only cell transcriptomes in which Td- tomato was detected. Cell types were identified by projecting cluster annotations from the main brain dataset onto *Cx3cr1*-Td+ dataset, as previously described (Stuart et al., 2019).

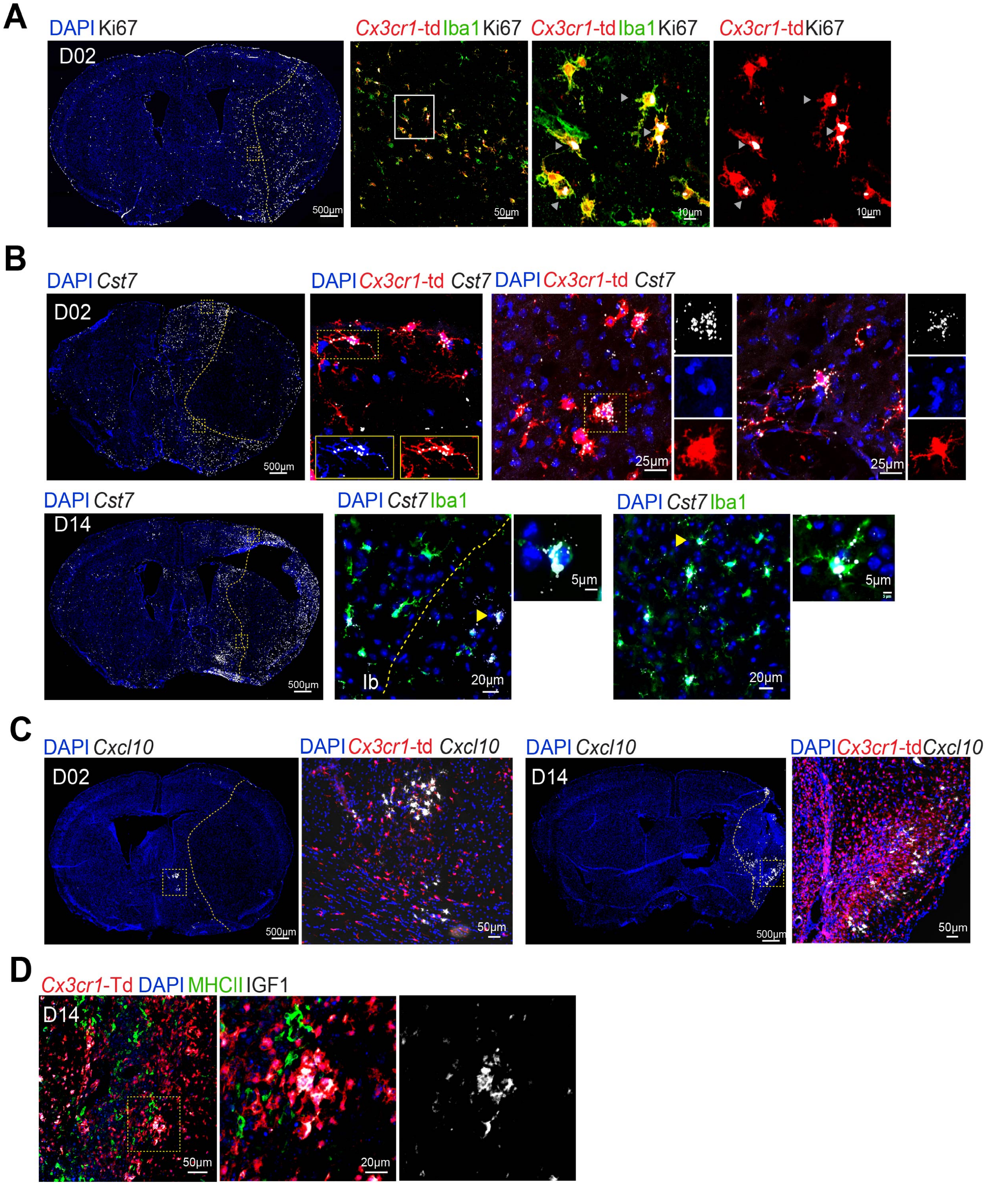


**Figure S6. Histological validation of microglia marker genes (Related to Figure 2H-K.)**

**(A)** Left: Representative immunofluorescence (IF) image of a whole brain section from a *Cx3cr1*-Td + mouse subjected to 2 days of MCAo (D02) showing the distribution of Ki67+ cells (white, binary mask) and nuclear DAPI staining (blue); Middle and right panels: IF images of magnified areas showing Ki67 expression by *Cx3cr1* -Td+(red) Iba1+(green) microglia in the peri-infarct area. Arrowheads indicate Ki67 staining. The border of the ischemic lesion is indicated by yellow dash outline and was traced based on DAPI, Iba1 and Td-tomato staining

**(B)** Top: RNAscope fluorescence in situ hybridization (FISH) combine with IF images validating *Cst7* (white) expression in D02 *Cx3cr1* -Td+ microglia (Td+, red). Left: Representative whole brain section image of *Cst7* expression (binary mask) and nuclear DAPI staining; Middle and right panels: FISH-IF images of magnified areas showing upregulation of *Cst7* in microglial cells surrounding the ischemic lesion. Bottom: FISH-IF images validating *Cst7* (white) expression in D14 mice. Left: Representative whole brain section image of *Cst7* expression (binary mask) and nuclear DAPI staining; Middle and right top panels: FISH-IF images of magnified areas showing upregulation of Cst7 in microglial cells (Iba1+, green) surrounding the ischemic lesion.

**(C)** FISH-IF images validating *Cxcl10* (white) expression in *Cx3cr1* -Td+ mice 2 and 14 days after MCAo. Left (D02): Representative whole brain section image of *Cxl10* expression (binary mask) and images of magnified areas showing localization of *Cxcl10* in microglial cells (Td+, red) outside of the ischemic lesion. Right (D14): Representative whole brain section image of *Cxl10* expression (binary mask) and images of magnified areas showing localization of*Cxcl10* in microglial cells (Td+, red) on the border of the ischemic lesion.

**(D)** IF images validating IGF1 (white) expression by *Cx3cr1* -Td+(red) MHCII– (green) microglia 14 days after MCAo in the ischemic region.

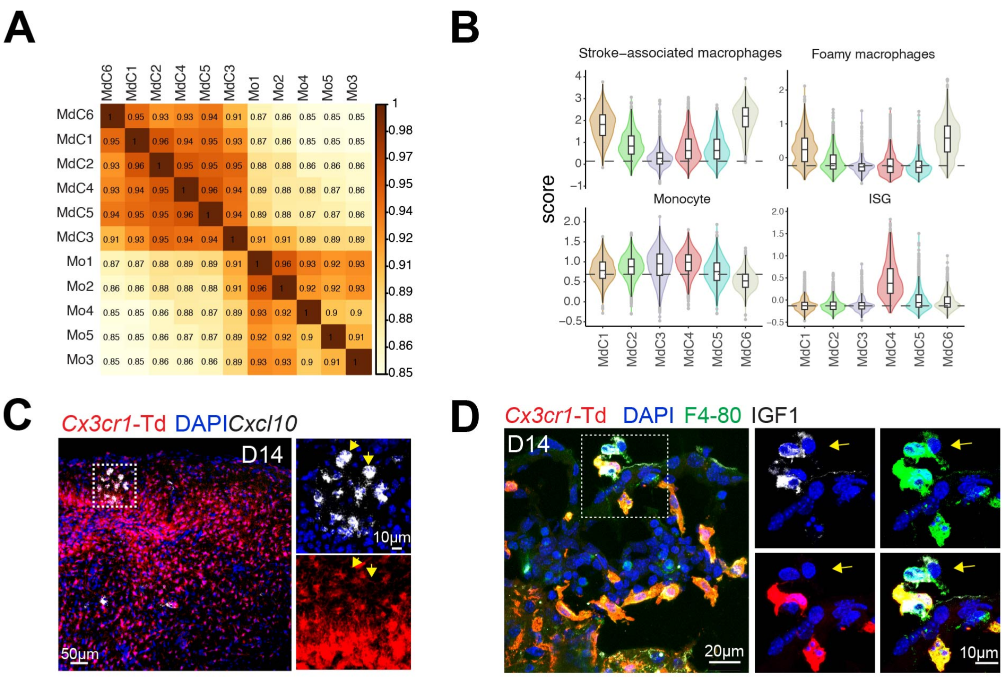


**Figure S7. Associative analysis, module scores and extended histology of blood monocytes and brain-infiltrated myeloid derived cells**

**(A)** Spearman correlation heatmap of gene expression average between peripheral blood monocytes (Mo) and brain infiltrating macrophages (MdC) clusters showing highest association of blood Mo clusters with MdC3 and the least association with MdC1 and MdC6. Scale bar represents Spearman correlation coefficient (r).

**(B)** Violin plots comparing gen module scores of brain infiltrating macrophage clusters (MdC) for stroke-associated macrophages, foamy macrophages, monocytic and interferon-stimulated genes (ISG) enrichment data sets. Genes used for calculating the module scores are listed in Table S4.

**(C)** FISH-IF images showing Cxcl10 (white) expression in D14 cells, which are localized near Cxcl10-expressing *Cx3cr1* -Td+ microglia (Td+, red). Nuclei are stained with DAPI (blue).

**(D)** IF images showing IGF1 (white) expression by surface *Cx3cr1* -Td– F4-80– (green) MdC,14 days after MCAo.

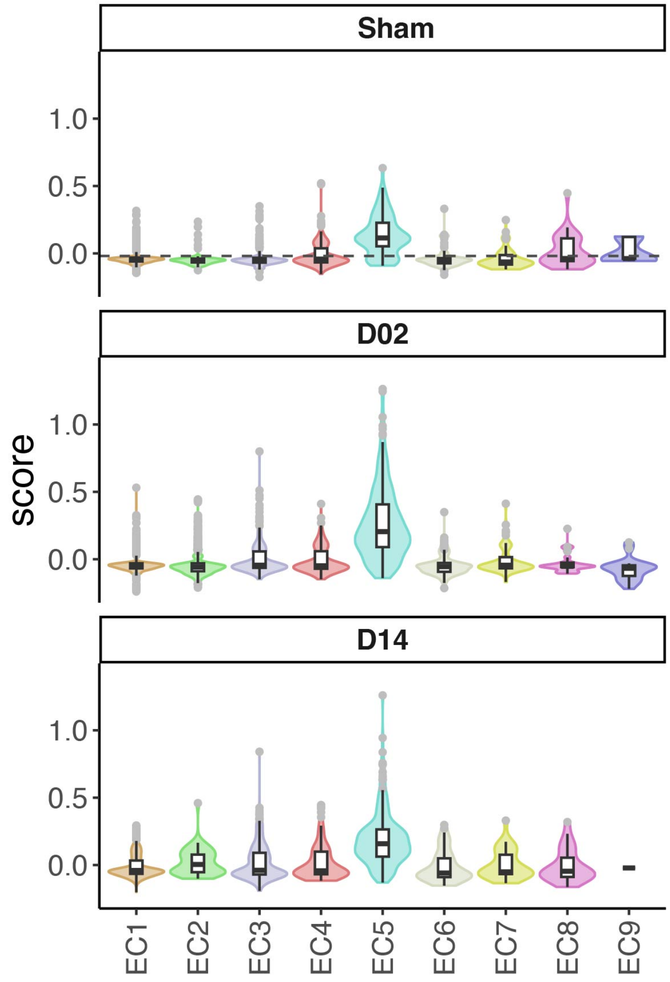


**Figure S8. ISG module score in endothelial cells**
Violin plots comparing module score distribution for interferon-stimulated genes (ISG) among brain endothelial cell clusters, stratified by control surgery (Sham), 2 days (D02) and 14 days (D14) stroke groups. Genes used for calculating the ISG module score are listed in Table S4.

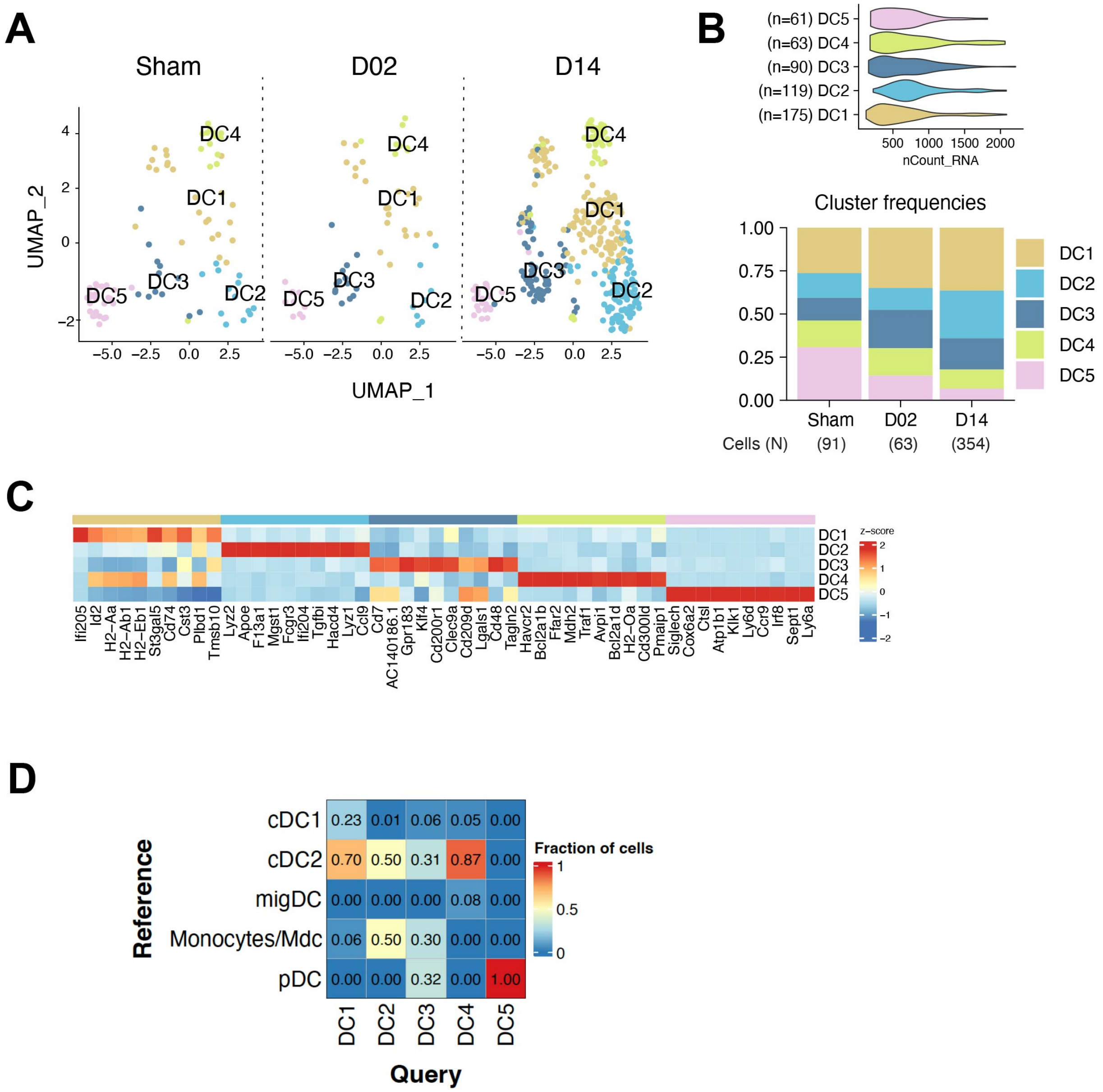

**Figure S9. Transcriptional changes in peripheral blood dendritic cells**

(A) UMAP plot of peripheral blood dendritic cells transcriptomes (DC) by studied time point: control surgery or stroke mice 2 or 14 days after injury (Sham, D02, D14, respectively).
(B) Top: Violin plot showing DC numbers in each cluster; Bottom: Bar graph showing relative frequencies of DC clusters across Sham, D02 and D14 groups.
(C) Heatmap displaying expression of the top 10 upregulated genes in each DC cluster. Scale bar represents Z-score of average gene expression (log).
(D) Heatmap of blood DC data sets showing the percentage of cell identity of each cluster (columns) attributed to a cell type classifier (rows) after unsupervised annotation with BrainImmuneAtlas reference dataset.

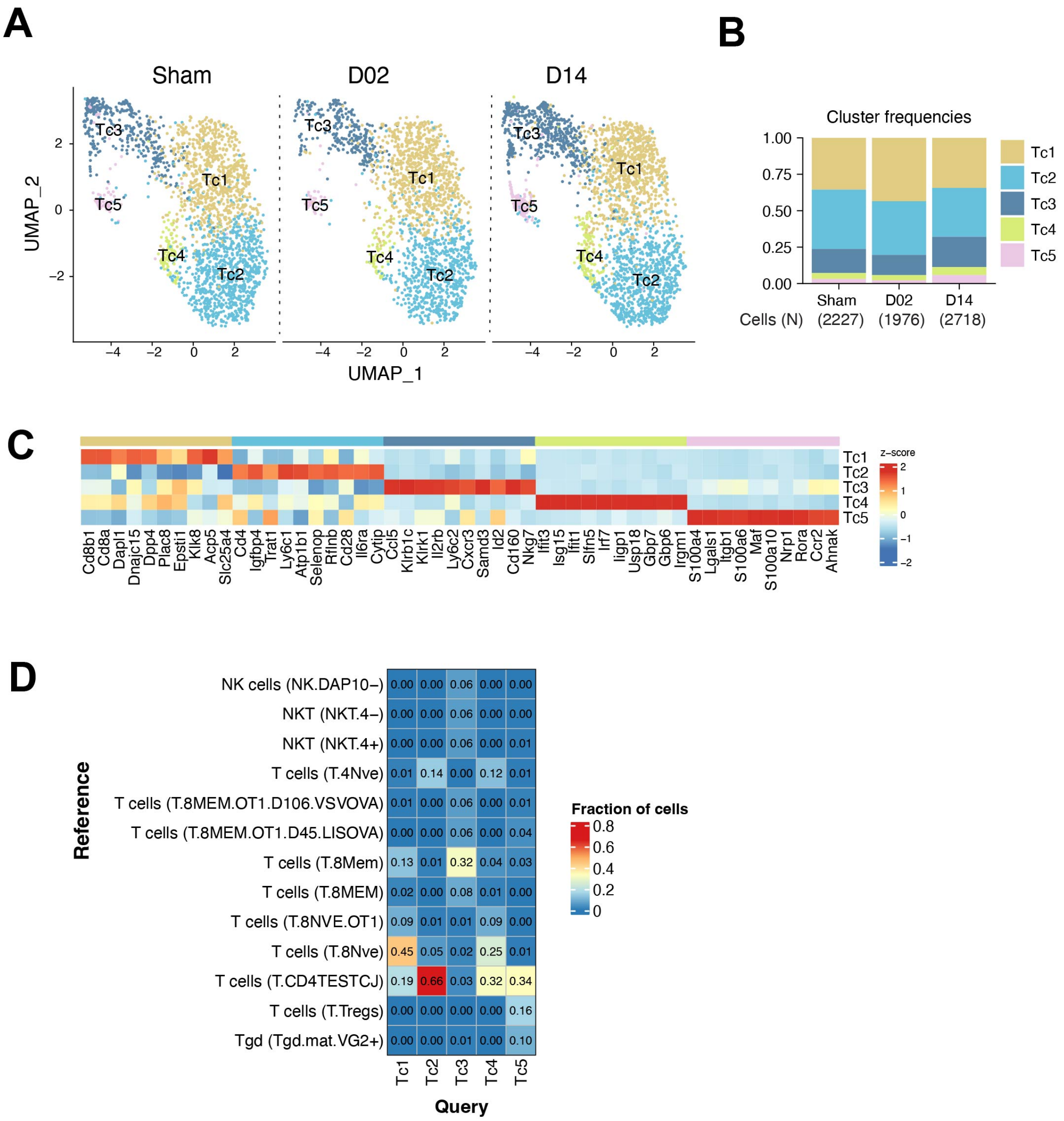


**Figure S10. Transcriptional changes in T cells from peripheral blood**

**(A)** UMAP plot of peripheral blood T cells transcriptomes (Tc) by studied time point: control surgery or stroke mice 2 or 14 days after injury (Sham, D02, D14, respectively).

**(B)** Bar graph showing relative frequencies and total cell numbers of Tc clusters across Sham, D02 and D14 groups.

**(C)** Heatmap displaying expression of the top 10 upregulated genes in each Tc cluster. Scale bar represents Z-score of average gene expression (log).

**(D)** Heatmap of blood Tc data sets showing the percentage of cell identity of each cluster (columns) attributed to a cell type classifier (rows) after unsupervised annotation with the ImmGen reference dataset.

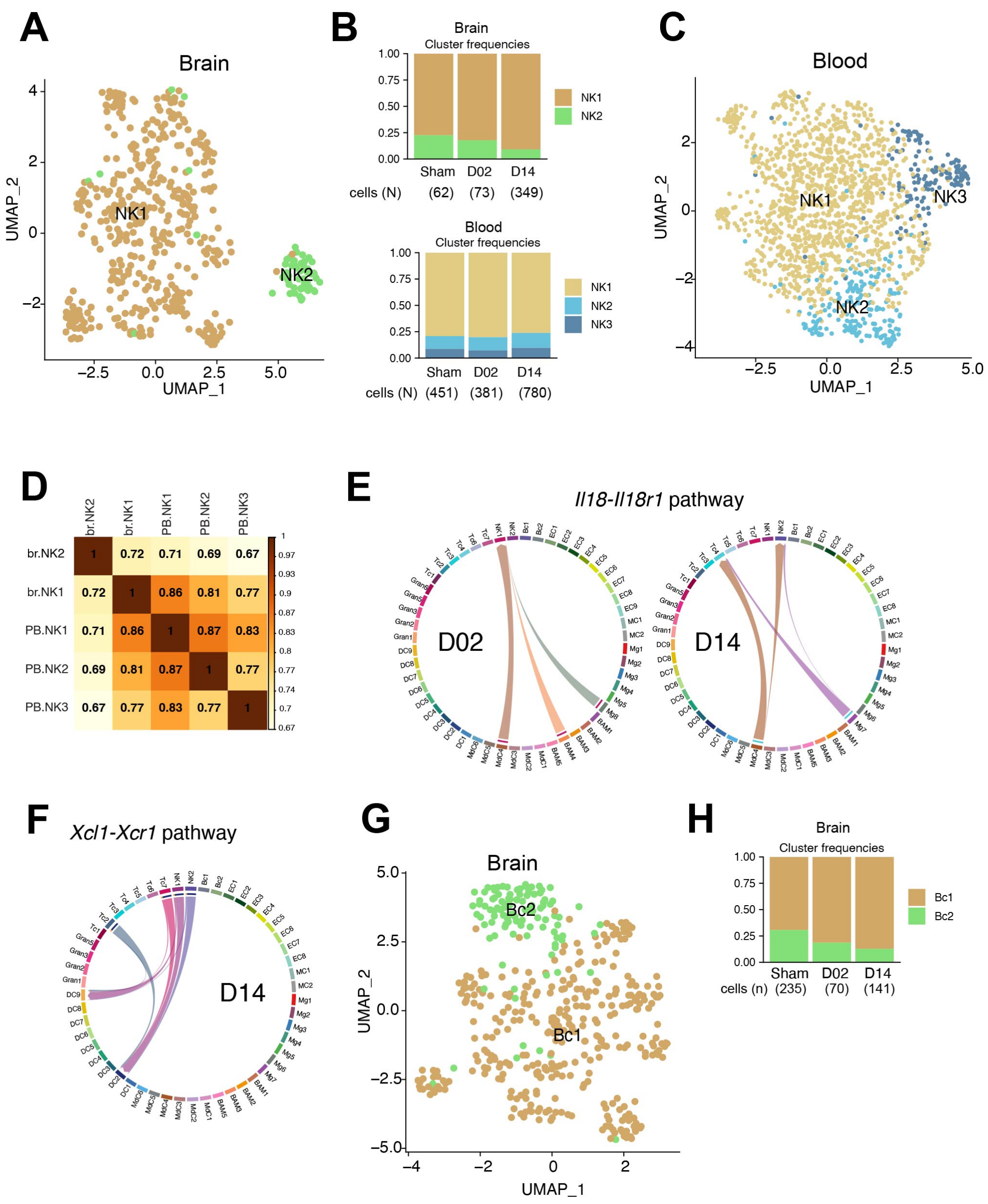

**Figure S11. Transcriptional changes in NK and B cells**

**(A)** UMAP plot of merged Sham, D02 and D14 brain NK transcriptomes identified 2 subclusters.

**(B)** Bar graph showing relative frequencies and total cell numbers of brain (Top) and blood (Bottom) NK clusters across Sham, D02 and D14 groups.

**(C)** UMAP plot of merged Sham, D02 and D14 blood NK transcriptomes identified 3 subclusters.

**(D)** Spearman correlation heatmap between blood (PB.NK) and brain (Br.NK) NK clusters showing high association of average gene expression of PB.NK clusters with Br.NK1 cluster. Scale bar represents Spearman correlation coefficient (r).

(**E,F**) Chord plots showing brain cell-cell interactions between Il18 and Il18r1 in D02 and D14 stroke mice (**E**) and *Xcl1-Xcr1* interaction in D14 mice (**F**). The strength of the interaction is indicated by the edge thickness. The color of the chord matches the cell cluster color sending the signal *(Il18* or *Xcl1* ). The number of cell cluster receptors *(Il18r1* or *Xcr1* ), and their weight in the interactions, is indicated by the color-matched stacked bar next to each cluster sender.

**(G)** UMAP plot of merged Sham, D02 and D14 brain B cells transcriptomes reveals 2 subclusters.

**(H)** Bar graph showing relative frequencies and total cell numbers of brain B cells clusters across Sham, D02 and D14 groups.

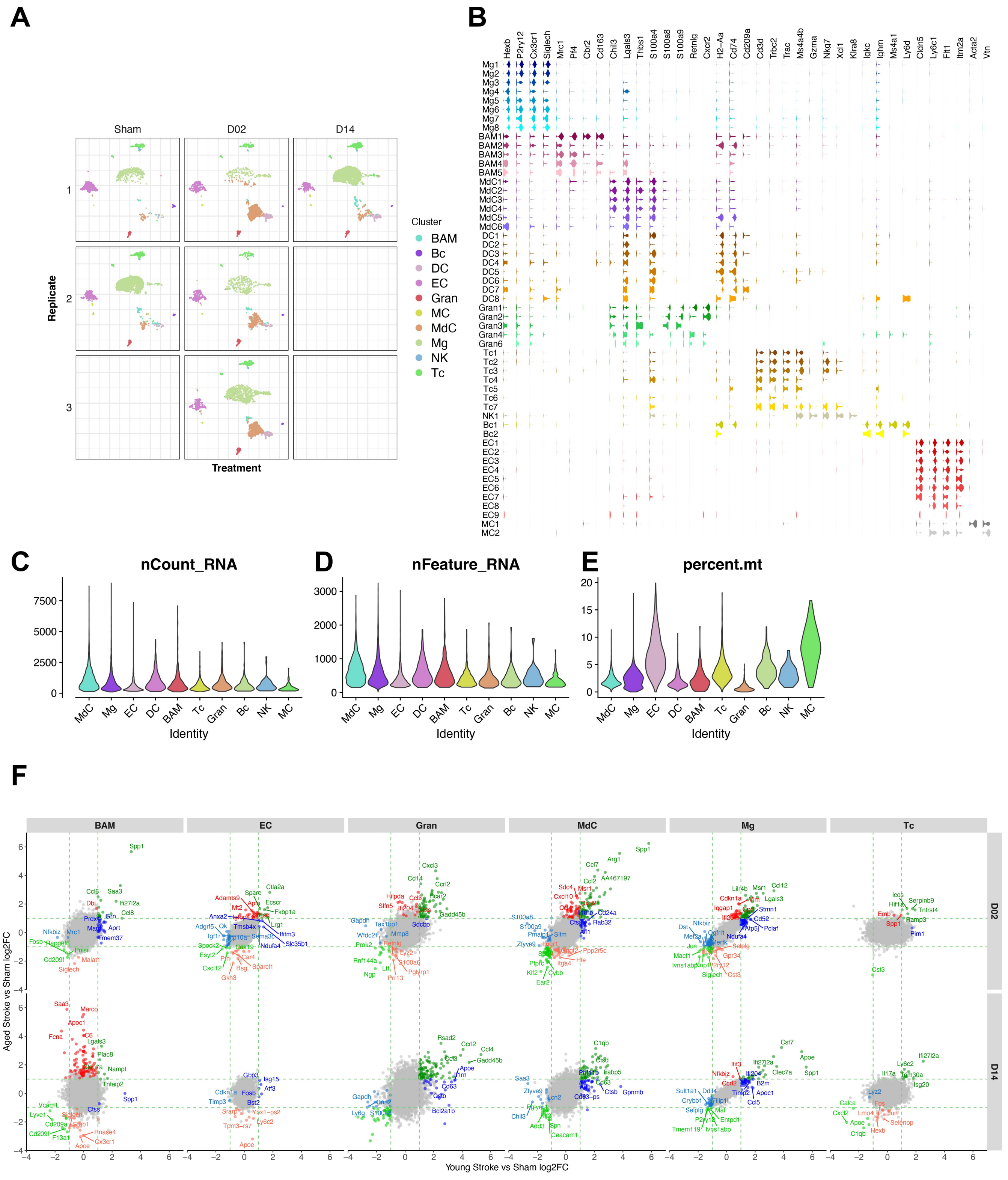


**Figure S12. Brain cell transcriptomic data set quality in aged mice and analysis of differential gene expression between aged and young stroke mice.**

**(A)** UMAPs of the identified clusters for each biological replicates of aged brain cell transcriptomes from control surgery (Sham) or stroke mice 2 (D02) or 14 (D14) days after injury.

(**B**) Violin plots generated from the integrated dataset displaying characteristic cell marker genes of each identified cell subcluster.

(**C, D, E)** Violin plots showing distribution of number of total UMI counts per cell (nCount) (**C**), genes detected per cell (nFeature) (**D**), and percentage of mitochondrial genes (percent.mt) per identified cell type (**E**). Microglia (Mg), border-associated macrophages (BAM), monocyte-derived cells (MdC), granulocytes (Gran), dendritic cells (DC), T cells (Tc), NK cells (NK), B cells (Bc), endothelial cells (EC), vascular mural cells (MC).(**F**) Analysis of differential gene expression between aged and young stroke mice. Scatterplots comparing stroke-induced differential gene expression for each brain cell type versus Sham in young and aged mice at D02 and D14. Genes with log2FC > 1 are highlighted in color. Green color indicates differentially regulated genes in both aged and young mice. Blue color indicates differentially regulated genes in only young mice. Red color indicates differentially regulated genes in only aged mice. Grey color indicates genes that were not found to be significantly changed in either aged or young stroke mice.
