## Supplementary material for "Brain and blood single-cell transcriptomics in acute and subacute phases after experimental stroke": Tables

**Table 1. Antibodies used for cytometric analysis and sorting**

| Antigen | Label | Clone | Isotype | Supplier | RRID |
|---|---|---|---|---|---|
| **CD11b** | APC/Cy7 | M1/70 | rat IgG2b, κ | Biolegend | AB_830641 |
| **CD11c** | PE/Cy7 | N418 | Armenian hamster | Biolegend | AB_493569 |
| **CD16/CD32** | | 93 | rat IgG2b, λ | Biolegend | AB_312800 |
| **CD172a** | AF700 | P89 | Rat IgG1, κ | Biolegend | AB_2650812 |
| **CD4** | F488 | RM4-5 | rat IgG2a, κ | Biolegend | AB_493373 |
| **CD45** | BV510 | 30F-11 | rat IgG2b, κ | Biolegend | AB_2561392 |
| **CD45** | APC | 30F-11 | rat IgG2b, κ | Biolegend | AB_312976 |
| **CX3CR1** | PE | SA011F11 | rat IgG2a, κ | Biolegend | AB_2564314 |
| **F4/80** | PE-Cy5 | BM8 | rat IgG2a, κ | Biolegend | AB_893494 |
| **I-A/I-E** | BV605 | M5/114.15.2 | rat IgG2b, κ | Biolegend | AB_2565894 |
| **Ly6C** | FITC | HK1.4 | rat IgG2c, κ | Biolegend | AB_1186134 |
| **Ly6G** | PerCP/Cy5.5 | 1A8 | rat IgG2a, κ | Biolegend | AB_1877272 |
| **Ly6G** | APC | 1A8 | rat IgG2a, κ | Biolegend | AB_1877163 |
| **TCR β** | PerCP/Cy5.5 | H57-597 | Armenian hamster | Biolegend | AB_1575176 |
| **TER-119** | Biotin | TER-119 | rat IgG2b, κ | Biolegend | AB_313704 |
| **XCR1** | PerCP/Cy5.5 | ZET | mouse IgG2b, κ | Biolegend | AB_2564363 |
| **CD197 (CCR7)** | PE | 4B12 | rat IgG2a, κ | Biolegend | AB_2564363 |
| **CD209a** | PE | MMD3 | mouse IgG2c, κ | Biolegend | AB_2721636 |
| **Lin:** | | | | | |
| **CD19** | FITC | 6D5 | rat IgG2a, κ | Biolegend | AB_313640 |
| **CD3ε** | FITC | 145-2C11 | Armenian hamster | Biolegend | AB_312670 |

| | | | | | |
|---|---|---|---|---|---|
| **Ly6G** | FITC | 1A8 | rat IgG2a, κ | Biolegend | AB_1236488 |
| **NK1.1** | FITC | PK136 | mouse IgG2a, κ | Biolegend | AB_313392 |
| **TCR β** | FITC | H57-597 | Armenian hamster | Biolegend | AB_313428 |

### Table S2. Drop-Seq run description

| Sample Name | Microfluidic Device | Bead Lot Number | Guanidine HCL USED? | Bead No. collected (theoretical ) | Bead No. (after Wash) | Bead No. (before cDNA Amp.) | Amp. Cycles | cDNA (ng/µl) | Total cDNA Yield ng | Library (ng/µl) |
|---|---|---|---|---|---|---|---|---|---|---|
| GR200817 | 12/18/19 0.9mm | 101619 | YES | 120,000 | 80,163 | 87,888 | 15 | 0.194 | 77.6 | 2.26 |
| GR210225 | 12/18/19 0.9mm | 101619 | YES | 93,333 | 78,107 | 65,690 | 15 | 0.0811 | 25.952 | |
| GR200812 | 12/18/19 0.9mm | 101619 | YES | 96,666 | 76,502 | 69,660 | 15 | 0.191 | 68.76 | 2.4 |
| GR200728 | 12/18/19 0.9mm | 101619 | YES | 113,333 | 102,773 | 74,894 | 15 | 0.172 | 67.08 | 1.7 |
| GR190610 | 10/04/18 0.9mm | 012819C | YES | 104,000 | 98,662 | 69,841 | 15 | 0.313 | 100.16 | 4.06 |
| GR200723 | 12/18/19 0.9mm | 101619 | YES | 53,333 | 51,386 | 39,161 | 15 | 0.202 | 40.4 | 2.8 |
| GR180426 | | 83117 | no | 125,000 | 78,800 | 50,780 | 14 | 0.524 | 125.76 | |
| GR180613tag | | 83117 | no | 152,000 | 102,773 | 64,607 | 15 | 0.305 | 122 | |
| GR180614tag | | 83117 | no | 164,000 | 131,550 | 82,835 | 15 | 0.175 | 70 | |
| GR180905TAG | | 83117 | YES | 112,000 | 53,442 | 48,546 | 15 | 0.428 | 102.72 | 2.1 |
| GR180919TAG | | 83117 | YES | 136,000 | 92,496 | 81,210 | 15 | 0.443 | 132.9 | 2.72 |
| GR190110 | 10/4/18 | 72818 | YES | 160,000 | 106,884 | 82,775 | 15 | 0.159 | 47.7 | 2.6 |
| GR180716TAG | | 83117 | no | 157,320 | 133,605 | 79,406 | 15 | 0.188 | 75.2 | 2.18 |
| GR181128 | 10/4/18 | 72818 | YES | 160,000 | 156,215 | 82,912 | 15 | 0.221 | 66.3 | 3.04 |
| GR181114 | 10/4/18 | 72818 | YES | 160,000 | 168,548 | 131,923 | 15 | 0.145 | 69.6 | 2.4 |
| GR181024 | 10/4/18 | 83117 | YES | 144,000 | 174,714 | 91,317 | 15 | 0.394 | 141.84 | 2.22 |
| GR181212 | 10/4/18 | 72818 | YES | 160,000 | 122,906 | 88,790 | 15 | 0.142 | 42.6 | 2.56 |
| GR180125 | | 83117 | no | 176,000 | 147,000 | 53,780 | 16 | | | |
| GR180517 | | 83117 | no | 150,000 | 96,600 | 83,196 | 16 | 0.213 | 85.2 | 2.372 |
| GR18110701 | 10/4/18 | 83117 | YES | 160,000 | 147,993 | 94,385 | 15 | 0.319 | 114.84 | 2.86 |
| GR18110702 | 10/4/18 | 83117 | YES | 160,000 | 137,716 | 100,702 | 15 | 0.283 | 101.88 | 2.64 |
| GR181120 | 10/4/18 | 72818 | YES | 160,000 | 121,272 | 136,795 | 15 | 0.143 | 68.64 | 3.46 |
| GR18121901 | 10/4/18 | 72818 | YES | 160,000 | 139,771 | 95,287 | 15 | 0.262 | 94.32 | 2.54 |
| GR18121902 | 10/4/18 | 72818 | YES | 160,000 | 129,494 | 85,421 | 15 | 0.147 | 44.1 | 2.18 |
| GR190116 | 10/4/18 | 72818 | YES | 160,000 | 168,548 | 91,678 | 15 | 0.285 | 98.325 | 2.44 |
| GR190410 | 10/04/18 0.9mm | 012819C | YES | 154,666 | 131,550 | 98,896 | 15 | 0.19 | 91.2 | 2.76 |
| GR190415 | 10/04/18 0.9mm | 012819C | YES | 160,000 | 152,104 | 118,868 | 15 | 0.148 | 82.88 | 2.22 |
| GR190417 | 10/04/18 0.9mm | 012819C | YES | 160,000 | 2,382 ( 103,3: | 74,714 | 15 | 0.309 | 30.9 | 2.68 |
| GR190522 | 10/04/18 0.9mm | 012819C | YES | 152,000 | 127,439 | 91,437 | 15 | 0.351 | 35.1 | 2.14 |
| GR190529 | 10/04/18 0.9mm | 012819C | YES | 160,000 | 131,550 | 90,234 | 15 | 0.174 | 83.52 | 2.56 |

| Sample Name | Mouse Strain | Sex | Age | No. of mice pooled | ECx10^3 | MGx10^3 | CD45hix10^3 | Td+cellsx10^3 | Cell populations within the sample | Treatment | Order Number | Index Primer | Illumina Run ID | Type (Next Seq/Miseq etc) | Sequencing sample ID | Seurat Designation |
|---|---|---|---|---|---|---|---|---|---|---|---|---|---|---|---|---|
| GR200817 | C57BL6_WT | female | 18m | 3 | 18 | 46 | 72 | N/A | EC (18K); MG(46K); CD45hi(72k); | D02 | | N701 | | | | |
| GR210225 | C57BL6_WT | female | 18m | 3 | 20 | 20 | 120 | N/A | EC (20K); MG(20K); CD45hi(120k); | D02 | | | | | | |
| GR200812 | C57BL6_WT | female | 18m | 2 | 25 | 124 | 54 | N/A | EC (25K); MG(124K); CD45hi(54k); | D14 | | N705 | | | | |
| GR200728 | C57BL6_WT | female | 18m | 4 | 32 | 141 | 44 | N/A | EC (32K); MG(141K); CD45hi(44k); | S02 | | N704 | | | | |
| GR190610 | CX3CR1_CreERhet_Tdhet | male | 8-12w | 4 | N/A | N/A | N/A | 125 | Td+cells ('MG";125K) | S02D02D14 | 10417514 | N704 | 190624_NS500503_0929_AH | NextSeq 500 | GR190610_GCTCAGGA | |
| GR200723 | C57BL6_WT | male | 17m | 2 | 12 | 25 | 18 | N/A | EC (12K); MG(25K); CD45hi(18k); | D02 | | N702 | | | | |
| GR180426 | C57BL6_WT | male | 8w | 6 | 24 | 24 | 200 | N/A | EC (24K); MG(24K); CD45hi(200k) | D02 | | | | | | D02R1 |
| GR180613tag | C57BL6_WT | male | 8w | 5 | 25 | 25 | 150 | N/A | EC (25K); MG(25K); CD45hi(150k); | D14 | 10400214 | N701 | 180705_NS500503_0712_AH | NextSeq 500 | GR180613TAG_TAAGGCGA | D14R2 |
| GR180614tag | C57BL6_WT | male | 8w | 5 | 25 | 25 | 150 | N/A | EC (25K); MG(25K); CD45hi(150k); | D02 | 10400214 | N702 | 180705_NS500503_0712_AH | NextSeq 500 | GR180614TAG_CGTACTAG | D02R2 |
| GR180905TAG | C57BL6_WT | male | 8w | 4 | 25 | 25 | 150 | N/A | EC (25K); MG(25K); CD45hi(150k); | D14 | 10404808 | N702 | 181012_NS500503_0767_AH | NextSeq 500 | GR180905TAG_CTAGTACG | D14R3 |
| GR180919TAG | C57BL6_WT | male | 8w | 5 | 25 | 25 | 150 | N/A | EC (25K); MG(25K); CD45hi(150k); | D02 | 10404808 | N704 | 181012_NS500503_0767_AH | NextSeq 500 | GR180919TAG_GCTCAGGA | D02R3 |
| GR190110 | C57BL6_WT | male | 8w | 4 | 33 | 134 | 33 | N/A | EC (33K); MG(134K); CD45hi(33k); | S02 | 10410132 | N704 | 190205_NS500503_0839_AH | NextSeq 500 | GR190110_GCTCAGGA | S02R4 |
| GR180716TAG | C57BL6_WT | male | 8w | 4 | 35.7 | 191 | 28.1 | N/A | EC (35.7K); MG(191K); CD45hi(28.1k); | S02 | 10404808 | N701 | 181012_NS500503_0767_AH | NextSeq 500 | GR180716TAG_TCGCCTTA | S02R1 |
| GR181128 | C57BL6_WT | male | 8w | 4 | 42 | 133 | 60 | N/A | EC (42K); MG(133K); CD45hi(60k); | S02 | 10408151 | N704 | 181213_NS500503_0808_AH | NextSeq 500 | GR181128_GCTCAGGA | S02R2 |
| GR181114 | C57BL6_WT | male | 9w | 4 | 45 | 125 | 209 | N/A | EC (45K); MG(125K); CD45hi(209k); | D14 | 10408151 | N702 | 181213_NS500503_0808_AH | NextSeq 500 | GR181114_CTAGTACG | D14R4 |
| GR181024 | C57BL6_WT | male | 9w | 5 | 50 | 50 | 100 | N/A | EC (50K); MG(50K); CD45hi(100k); | D02 | 10408151 | N701 | 181213_NS500503_0808_AH | NextSeq 500 | GR181024_TCGCCTTA | D02R4 |
| GR181212 | C57BL6_WT | male | 10w | 4 | 60 | 120 | 60 | N/A | EC (60K); MG(120K); CD45hi(60k); | S02 | 10410132 | N701 | 190205_NS500503_0839_AH | NextSeq 500 | GR181212_TCGCCTTA | S02R3 |
| GR180125 | C57BL6_WT | male | 8w | 5 | 43.2 | 47.3 | 109.5 | N/A | EC(43.2K); PMN(47.3K); Macro(109.5K) | D14 | 10391620 | | 180213_NB551191_0108_AH | NextSeq 500 | GR180125 | D14R1 |
| GR180517 | C57BL6_WT | male | 8w | 1 | N/A | N/A | N/A | N/A | Peripheral blood leukocytes (240K) | N | | | | | | |
| GR18110701 | C57BL6_WT | male | 9w | 2 | N/A | N/A | N/A | N/A | Peripheral blood leukocytes (240K) | S02 | 10407791 | N701 | 181211_NS500503_0806_AH | NextSeq 500 | GR18110701_TCGCCTTA | PBS02R1 |
| GR18110702 | C57BL6_WT | male | 9w | 2 | N/A | N/A | N/A | N/A | Peripheral blood leukocytes (240K) | D02 | 10407791 | N702 | 181211_NS500503_0806_AH | NextSeq 500 | GR18110702_CTAGTACG | PBD02R1 |
| GR181120 | C57BL6_WT | male | 9w | 2 | N/A | N/A | N/A | N/A | Peripheral blood leukocytes (240K) | D14 | 10407791 | N704 | 181211_NS500503_0806_AH | NextSeq 500 | GR181120_GCTCAGGA | PBD14R1 |
| GR18121901 | C57BL6_WT | male | 10w | 2 | N/A | N/A | N/A | N/A | Peripheral blood leukocytes (240K) | S02 | 10410132 | N702 | 190205_NS500503_0839_AH | NextSeq 500 | GR18121901_CTAGTACG | PBS02R2 |
| GR18121902 | C57BL6_WT | male | 10w | 2 | N/A | N/A | N/A | N/A | Peripheral blood leukocytes (240K) | D02 | 10414364 | N701 | 190426_NS500503_0890_AH | NextSeq 500 | GR18121902_TCGCCTTA | PBD02R2 |
| GR190116 | C57BL6_WT | male | 10w | 2 | N/A | N/A | N/A | N/A | Peripheral blood leukocytes (240K) | D02 | 10414364 | N702 | 190426_NS500503_0890_AH | NextSeq 500 | GR190116_CTAGTACG | PBD02R3 |
| GR190410 | C57BL6_WT | male | 9w | 2 | N/A | N/A | N/A | N/A | Peripheral blood leukocytes (240K) | D02 | 10416797 | N701 | 190611_NS500503_0919_AH | NextSeq 500 | GR190410_TCGCCTTA | |
| GR190415 | C57BL6_WT | male | 9w | 2 | N/A | N/A | N/A | N/A | Peripheral blood leukocytes (240K) | D14 | 10416797 | N702 | 190611_NS500503_0919_AH | NextSeq 500 | GR190415_CTAGTACG | |
| GR190417 | C57BL6_WT | male | 9w | 2 | N/A | N/A | N/A | N/A | Peripheral blood leukocytes (240K) | S02 | 10416797 | N704 | 190611_NS500503_0919_AH | NextSeq 500 | GR190417_GCTCAGGA | |
| GR190522 | C57BL6_WT | male | 8w | 2 | N/A | N/A | N/A | N/A | Peripheral blood leukocytes (240K) | S02 | 10417514 | N701 | 190624_NS500503_0929_AH | NextSeq 500 | GR190522_TCGCCTTA | |
| GR190529 | C57BL6_WT | male | 8w | 2 | N/A | N/A | N/A | N/A | Peripheral blood leukocytes (240K) | D14 | 10417514 | N702 | 190624_NS500503_0929_AH | NextSeq 500 | GR190529_CTAGTACG | |

**Table S3. Cell numbers among cell type classes of down-sampled young and aged brain datasets**

| | celltype | Aged | Young |
|---|---|---|---|
| 1 | Mg | 5970 | 5241 |
| 2 | BAM | 295 | 316 |
| 3 | MdC | 2112 | 2875 |
| 4 | DC | 470 | 921 |
| 5 | Gran | 301 | 324 |
| 6 | Tc | 836 | 380 |
| 7 | NK | 44 | 125 |
| 8 | Bc | 96 | 115 |
| 9 | MaC | 0 | 10 |
| 10 | EC | 1747 | 1611 |
| 11 | MC | 92 | 36 |
| 12 | Epi | 20 | 27 |
| 13 | OD | 16 | 18 |

#### Table S4. Gene lists of module scores

| DAM | DIM | ISG | SAM | Foamy macrophages | monocytes |
|---|---|---|---|---|---|
| Dkk2 | Atf3 | Rsad2 | Spp1 | Fabp4 | S100a4 |
| Fabp5 | Btg2 | Socs1 | Fabp5 | Ctsl | Ms4a6c |
| Gm1673 | Ccl4 | Ddx60 | Gpnmb | Atp6v0d2 | Crip1 |
| Gpnmb | Ctss | Ifit2 | Ctsb | Gpnmb | Ctss |
| Igf1 | Dusp1 | Ifit3 | Ctsl | Fabp5 | Ccl9 |
| Itgax | Egr1 | Ifit3b | Lgals3 | Htra1 | F13a1 |
| Mamdc2 | Fos | Herc6 | Lpl | Epb41l3 | Plac8 |
| Spp1 | Icam1 | Rtp4 | Fth1 | Pld3 | Fn1 |
| | Ier2 | Spats2l | Cd63 | | Ccr2 |
| | Ier5 | Ifi44l | Ctsd | | Psap |
| | Il1a | Oas3 | | | Ms4a4c |
| | Il1b | Cxcl10 | | | Npc2 |
| | Itga6 | Mx1 | | | Ifi30 |
| | Jun | Mx2 | | | Lyz2 |
| | Junb | Isg15 | | | Pld4 |
| | Klf6 | Oasl1 | | | Lamp1 |
| | Tnf | Ifi44 | | | Ifitm3 |
| | Zfp36 | Oas2 | | | Smpdl3a |
| | Cd83 | Usp18 | | | Ly86 |
| | Fosb | Epsti1 | | | Ctsc |
| | Cd14 | Oas1e | | | Vim |
| | Fth1 | Oas1c | | | Lgals1 |
| | Nfkbia | Oas1b | | | Clec4a3 |
| | Nfkbiz | Oas1f | | | S100a10 |
| | | Oas1h | | | Prdx1 |
| | | Oas1g | | | Anxa5 |
| | | Oas1a | | | Ctsb |
| | | Oas1d | | | Dbi |
| | | Siglec1 | | | Napsa |

| Module | Reference |
|---|---|
| DAM | Silvin et al. Immunity 2022 Aug 9;55(8):1448-1465.e6. doi: 10.1016/j.immuni.2022.07.004. |
| DIM | Silvin et al. Immunity 2022 Aug 9;55(8):1448-1465.e6. doi: 10.1016/j.immuni.2022.07.004. |
| ISG | Kim et al. Interferon Cytokine Res. 2018 Apr;38(4):171-185. doi: 10.1089/jir.2017.0127 |
| SAM | Beuker et al. Nat. Commun. 2022 Feb 17;13(1):945. doi: 10.1038/s41467-022-28593-1. |
| Foamy macr | Williams et al. Nat Immunol. 2020 Oct;21(10):1194-1204. doi: 10.1038/s41590-020-0768-4. |
| monocytes | Ballesteros et al. Cell 2020 Nov 25;183(5):1282-1297.e18. doi: 10.1016/j.cell.2020.10.003 |

**Table S5. Antibodies used for immunohistochemistry**

| Antigen | Biological source | Clone | Isotype | Supplier | Cat. Num. | RRID | Dilution |
|---|---|---|---|---|---|---|---|
| **IGF1** | mouse | Sm1.2 | IgG1k | Millipore | 05-172 | AB_309643 | 1:100 |
| **CD206** | rat | MR5D3 | IgG2a | BioRad | MCA2235 | AB_324622 | 1:200 |
| **Iba1** | rabbit | polyclonal | | Wako | 019-19741 | AB_839504 | 1:200 |
| **DsRed (TdTomato)** | rabbit | polyclonal | | Takara | 632496 | AB_10013483 | 1:200 |
| **Anti-MHC II (I-a/I-E)** | rat | M5/114 | IgG2bk | Millipore | MABF33 | AB_10807702 | 1:200 |

**Table S6. RNA scope probe list**

| | Reference | Name |
|---|---|---|
| 1 | 437511 | RNAscope Probe - Mm-Mrc1 |
| 2 | 480311-C2 | RNAscope Probe - Mm-Cd209a-C2 |
| 3 | 423381 | RNAscope Probe - Mm-Lrg1 |
| 4 | 426371-C2 | RNAscope Probe - Mm-Ms4a4c-C2 |
| 5 | 498711 | RNAscope Probe - Mm-Cst7 |
| 6 | 432871 | RNAscope Probe - Mm-Ccr7 |
| 7 | 403431-C3 | RNAscope Probe - Mm-Arg1-C3 |
| 8 | 408921-C2 | RNAscope Probe - Mm-Cxcl10-C2 |
| 9 | 546211-C2 | RNAscope Probe - Mm-Ccl8-C2 |
| 10 | 468421-C2 | RNAscope® Probe- Mm-Car4-C2 |
